# Inhibition of Endocannabinoid Degradation in Astrocytes Reprograms Glial Reactivity and Prevents Seizure Sequelae

**DOI:** 10.64898/2026.02.04.703855

**Authors:** Chudai Zeng, Fei Gao, Jian Zhang, Mei Hu, Dexiao Zhu, Li Sun, Jianlu Lyu, Mingzhe Pan, Chu Chen

## Abstract

**Background:** Temporal lobe epilepsy (TLE) is the most common form of focal epilepsy and is characterized by a pathological cascade of excitotoxicity that leads to neuroinflammation, progressive neuronal loss, and subsequent cognitive decline. Despite its prevalence, effective disease-modifying therapies remain lacking. Previous studies have demonstrated that the endocannabinoid system contributes to epileptic activity. In particular, inactivation of monoacylglycerol lipase (MAGL), the key rate-limiting enzyme responsible for the degradation of the endocannabinoid 2-arachidonoylglycerol (2-AG), an endogenous lipid mediator with anti-inflammatory and neuroprotective properties, suppresses seizures and reduces neuroinflammation. However, the cellular and molecular mechanisms underlying these protective effects remain unclear.

**Methods:** To dissect the cellular mechanisms underlying MAGL-mediated neuroprotection, we employed a kainic acid (KA)-induced status epilepticus model in mice with global, astrocyte-specific (aKO), and neuron-specific (nKO) deletion of *mgll*. We combined single-nucleus RNA sequencing (snRNA-seq) to map the transcriptomic landscape of glial responses with pharmacological interventions to validate key signaling pathways, as well as behavioral assays to assess functional recovery.

**Results:** We demonstrated that astrocyte-specific, but not neuron-specific, *mgll* deletion was sufficient to attenuate seizure susceptibility and hippocampal neurodegeneration, thereby recapitulating the protective phenotype observed in global knockouts. Transcriptomic profiling revealed that astrocytic MAGL deficiency fundamentally reshaped the glial response to injury by preventing the transition to pro-inflammatory reactive astrocyte states and suppressing the activation of disease-associated microglia (DAM). Mechanistically, we identified a signaling pathway in which the neuroprotective effects of MAGL inhibition depend on cannabinoid receptor 1 (CB1) activation and are mediated by downstream peroxisome proliferator-activated receptor γ (PPAR-γ) signaling. Either genetic deletion of CB1 or pharmacological blockade of PPAR-γ abolished the protective effects. Furthermore, aKO mice exhibited reduced neuronal loss, preserved synaptic structural integrity and protection against post-seizure cognitive deficits.

**Conclusion:** These findings reveal astrocytic MAGL as a crucial regulatory node in the epileptic brain and demonstrated that enhancing 2-AG signaling in astrocytes orchestrates neuroprotection via CB1-PPAR-γ signaling pathways, thereby attenuating neuroinflammation, preserving synaptic function, and preventing the cognitive comorbidities associated with epilepsy.

## Introduction

Temporal lobe epilepsy (TLE), particularly acquired forms, is characterized not only by recurrent spontaneous seizures but also by a devastating sequence of events often initiated by a precipitating insult, such as status epilepticus (SE) [1]. This initial insult triggers acute excitotoxicity, leading to a cascade of neuroinflammation and progressive neuronal loss [2]. Although many antiseizure medications are available, nearly 30∼40 % of patients do not respond to common drugs, leading to drug-resistant epilepsy. While current therapies often target neuronal excitability, they fail to address the underlying mechanisms that drive seizure recurrence and disease progression[3]. In particular, inflammatory responses orchestrated by non-neuronal cells are increasingly viewed as a pivotal determinant of the long-term fate of the injured brain. Understanding how to modulate these glial-driven inflammatory cascades represents a promising avenue for neuroprotective intervention during the epileptogenic window. The endocannabinoid (eCB) system has emerged as a potent brake on this pathological progression, integrating synaptic activity with inflammatory and metabolic signaling. This neuromodulatory system comprises endogenous cannabinoids (primarily 2-arachidonoylglycerol [2-AG] and anandamide [AEA]), cannabinoid receptors (CB1 and CB2), and the enzymes responsible for endocannabinoid synthesis and degradation [4, 5]. Upon neuronal depolarization and calcium influx, 2-AG, the most abundant endocannabinoid in the brain[5], is synthesized on-demand from membrane phospholipids by diacylglycerol lipase-α (DAGLα), released into the synaptic cleft, and travels retrogradely to activate presynaptic cannabinoid type 1 receptors (CB1R), thereby reducing glutamate release and providing a critical negative feedback mechanism against hyperexcitability [6]. In experimental epilepsy models, eCB signaling has been implicated in limiting excitotoxicity and dampening network hyperactivity [7, 8]. Moreover, clinical success of cannabidiol in treating drug-resistant epilepsies has renewed interest in targeting the eCB system for epilepsy treatment [9]. Critically, the eCB system serves as a critical interface between neuronal activity and inflammatory signaling. While 2-AG signaling can exerts broad anti-inflammatory effects through CB1R, its hydrolysis releases arachidonic acid (AA), a precursor for cyclooxygenase (COX)-mediated production of proinflammatory prostaglandins (particularly PGE2) [5, 10–12]. This dual nature of 2-AG metabolism underscores its complex involvement in both seizure control and neuroinflammation.

Monoacylglycerol lipase (MAGL; gene name: *mgll*) is the primary enzyme responsible for degrading 2-arachidonoylglycerol (2-AG) in the brain. In various epilepsy models, global inhibition of *mgll*, which enhances 2-AG signaling while reducing its downstream metabolites, has been shown to suppress seizure activity [7, 13–15] and reduce neuroinflammation [15]. These findings collectively establish the therapeutic potential of targeting MAGL to enhance endocannabinoid tone. However, these systemic approaches cannot distinguish the cell type-specific roles of 2-AG metabolism in attenuating epileptic activity, leaving a critical gap in our mechanistic understanding and limiting the development of targeted, precision therapeutic strategies.

In this study, we dissected the cell-specific role of 2-AG signaling in altering the trajectory of KA-induced post-seizure injury. We report that astrocytic MAGL deletion suppresses neuroinflammation in the subacute phase and provides sustained protection into the chronic phase, whereas neuronal deletion offers no such benefit. Mechanistically, these protective effects are mediated via CB1 and PPARγ signaling pathways, establishing astrocytic metabolic reprogramming as a key strategy to suppress post-seizure injury.

## Materials and methods

### Animal

Mgll^flox/flox^ animals were generated by the Texas A&M Institute for Genomic Medicine. (TIGM) using a targeting strategy, as described previously[16]. Briefly, the targeting vector was introduced into the C57BL/6N ES cell line (JM8) via homologous recombination to flank Exon 2 with *LoxP* sites. To validate the conditional strategy, global *mgll* knockout (MAGL KO) mice were generated by crossing mgll^flox/flox^ mice with the Sox2-Cre line (Tg(Sox2-cre)1Amc/J, JAX #004783). Correct targeting and recombination were verified using long-distance PCR and sequencing. Cell-type-specific knockouts were generated as follows: Neuronal (nKO): mgll^flox/flox^ crossed with Syn1-Cre (JAX #003966). Astrocytic (aKO): mgll^flox/flox^ crossed with GFAP-Cre (JAX #024098). For mechanistic validation, global Cannabinoid Receptor 1 knockout mice (Cnr1^−/-^, CB1-KO) were also utilized. Experimental cohorts included both male and female mice (10∼12 weeks of age). Animals from each genotype were randomly assigned to either vehicle or kainic acid (KA) treatment groups. All mice were housed in a temperature-controlled facility on a 12-hour light/dark cycle with ad libitum access to food and water. All animal studies were performed in compliance with the US Department of Health and Human Services Guide for the Care and Use of Laboratory Animals, and the care and use of the animals reported in this study were approved by the Institutional Animal Care and Use Committee of University of Texas Health San Antonio.

### Kainic Acid-Induced Status Epilepticus Model

To induce Status Epilepticus (SE), mice received a single intraperitoneal (*i.p.*) injection of 17.5 mg/kg Kainic Acid (KA; HB0355, Hellobio) dissolved in sterile saline. Control mice received an equivalent volume of saline vehicle. Following injection, mice were monitored continuously for 2 hours to evaluate seizure severity using the modified Racine scale: Stage 1, immobility and staring; Stage 2, head nodding; Stage 3, unilateral forelimb clonus; Stage 4, bilateral forelimb clonus and rearing; Stage 5, rearing and falling; Stage 6, generalized tonic-clonic seizures with falling; Stage 7, seizures leading to death. Seizure incidence was defined as the occurrence of at least one seizure of Stage 4 or higher.

### Pharmacological Interventions

For pharmacological inhibition of MAGL, CB1R knockout mice were intraperitoneally injected with the selective inhibitor 4-Nitrophenyl-4-[bis (1,3-benzodioxol-5-yl)(hydroxy)methyl]piperidine-1-carboxylate (JZL184, 10 mg/kg) or vehicle (DMSO) was administered every other day for 3 injections. JZL184 was prepared and dissolved in a vehicle containing Tween 80 (10%), DMSO (10%) and saline (80%) as described previously [17]. To investigate the role of PPAR-γ, the antagonist GW9662 (5 mg/kg, *i.p.*; Cayman Chemical) was administered 30 minutes prior to JZL-184 treatment [18]. Mice were randomly assigned to treatment groups and monitored for seizure susceptibility as described above.

### Behavioral Assays

#### Open Field Test (OFT)

Hyperactive and anxiety-like behavior were assessed in an open field apparatus (30 × 30 cm) as described previously [16, 19]. Mice were placed in the center of the arena and allowed to explore freely for 30 minutes. Movement was tracked using an automated video tracking system (EthoVision, Noldus, version 17). Total distance travelled, mean activity, and time spent freezing were recorded.

#### Novel Object Recognition (NOR)

The NOR test was conducted in a 30 × 30 × 30 cm square open field as described previously [16, 20]. Briefly, the NOR test consisted of three phases: habituation, training, and testing. Mice first underwent a 10-minute habituation session in the empty arena. The NOR assay then comprised two 10-minute stages separated by a 4-hour intertrial interval. In the training stage, animals were presented with two identical objects. In the test stage, animals were presented with one familiar object (from training) and one novel object. Exploration was defined as the frequency with the head oriented toward and within 2 cm of an object. Object exploration during the test stage was quantified using the EthoVision video-tracking system (Noldus, version 17). The recognition index (RI) was calculated based on the following equation: RI =FN/FN+FF, where FN is the frequency devoted to the novel object and FF is the frequency for the familiar object.

### Tissue Processing and Immunofluorescence

Animals were anesthetized with ketamine/xylazine (200/10 mg/kg) and transcardially perfused with ice-cold PBS followed by 4% paraformaldehyde (PFA), as described previously [16, 21, 22]. Brains were post-fixed, cryoprotected in 30% sucrose, and sectioned at 25 μm using a cryostat. Free-floating sections were blocked with 5% goat serum and incubated overnight at 4°C with the following primary antibodies: anti-GFAP (1:1000, Cat# G3893, MilliporeSigma), anti-Iba1 (1:1000, Cat# MABN92, MilliporeSigma), and anti-PSD-95 (1:500, Cat# Ab2723, Abcam). Sections were then incubated with fluorophore-conjugated secondary antibodies followed by incubation with the corresponding fluorescent-labeled secondary antibody. 4-6-Diamidino-2-phenylindole (DAPI). Images were acquired using a Zeiss Imager II deconvolution microscope with SlideBook 2024 software.

### Histochemistry

Degenerated neurons were detected using Fluoro-Jade C (FJC), which is an anionic dye that specifically stains the soma and neurites of degenerating neurons and thus is unique as a neurodegenerative marker. As described previously [16, 22, 23], cryostat cut sections were incubated in the solution with FJC (0.0001% solution, Cat# TR-160-FJC, Biosensis) and DAPI (0.5 μg/ml) for 10 min, followed by 3× 1-min wash with distilled water. Slices were dried naturally at room temperature without light. The FJC positive cells in the cortex and hippocampus were imaged and analyzed using a Zeiss deconvolution microscope with SlideBook 2024 software.

### Quantitative Real-Time PCR (qPCR)

Total RNA was extracted from harvested tissues with the RNeasy Mini Kit (Qiagen) and treated with RNase-free DNase (Qiagen), following the manufacturer’s instructions. The RNA concentration was measured by a Microvolume UV-Vis Spectrophotometer (NanoDrop One, ThermoFisher Scientific), and RNA integrity was verified by electrophoresis in a 1% agarose gel. Reverse transcription was performed using the iScript cDNA synthesis kit (BioRad) with 1 µg of total RNA, with 4 µl of 5× iscript reaction mix, and 1 µl of iscript reverse transcriptase in a final volume of 20 µl. Samples were incubated at 25 °C for 5 min, followed by 42 °C for 30 min, and reactions were stopped by heating to 85 °C for 5 min. Real-time RT-PCR primers specific for IL-1β, IL-6, TNFα, and GAPDH were designed using Beacon Designer Software (BioRad) and synthesized by IDT (Coralville, IA). Primer sequences are listed in Table S1.

Reactions were performed in duplicate in a 25 µL total volume containing 12.5 µL of 2× iQ SYBR Green Supermix (Bio-Rad), 5 µL of 1:10 diluted cDNA template, and 400 nM of each primer. The PCR cycling protocol was as follows: 95 °C for 3 minutes, followed by 45 cycles of 95 °C for 30 seconds, 58 °C for 45 seconds, and 95 °C for 1 minute. A melt-curve analysis was conducted at the end of each run to confirm amplification specificity. Amplicon sizes were further verified by electrophoresis on a 3% agarose gel. Amplification and analysis were performed using the iCycler iQ Multicolor Real-Time PCR Detection System (Bio-Rad). Relative gene expression was quantified using the 2-^ΔΔCT^ method, with β-actin as the internal control as its expression was not affected by our experimental treatments. ΔCT was calculated as (CT of gene of interest) - (CT of β-actin), and ΔΔCT as (ΔCT of treated sample) - (ΔCT of control), as previously described [16, 21, 22, 24, 25]. Technical replicates were performed for each detected gene to ensure measurement accuracy and reproducibility.

### Public Bulk RNA sequencing Dataset Analysis

To evaluate the endogenous transcriptional dynamics of *mgll* relative to neuroinflammation, we re-analyzed publicly available bulk RNA-sequencing datasets obtained from the NCBI Gene Expression Omnibus (GEO). We utilized dataset GSE213393, comprising transcriptomic profiles from the mouse hippocampal dentate gyrus (DG) following kainic acid (KA) or saline injection for 3, 7, and 14 days. Additionally, dataset GSE99577 was employed to examine temporal gene expression changes in the CA1 region across a time course of 6 hours to 12 days post-injection. Normalized expression data were retrieved and processed using R. For robust comparison across samples, raw expression values for each gene were Z-score normalized. To quantify neuroinflammation, a composite neuroinflammation score was derived by calculating the mean of the Z-normalized expression values for a defined panel of canonical astrocytic markers, microglial markers, cytokines, and complement system. Temporal dynamics and correlations between *mgll* and this inflammation score were visualized using time-course line plots and heatmaps.

### Single-nucleus RNA Sequencing

#### Sample preparation, library construction, and RNA sequencing

Hippocampal tissues from WT and aKO mice treated with Veh or KA for single-nucleus RNA sequencing were harvested and stored in liquid nitrogen as previously described [26–28]. Single nucleus sample preparation, library construction, and RNA sequencing were outsourced to Novogene Corporation Inc. (Sacramento, CA). Single nucleus RNA sequencing libraries were prepared using the Chromium Next GEM Single Cell 3′ Gel Bead v4 Kit (10x Genomics) and sequencing was performed on the Illumina NovaSeq6000 system at Novogene.

#### RNA-Seq Data Processing and Analysis

Sequencing data were processed using CellBender (v4.0) to remove ambient RNA profiles. Initial filtering was performed using emptyDrops to retain droplets with an FDR < 0.01. Strict quality control was applied to remove low-quality nuclei: cells were excluded if they exhibited >15% mitochondrial reads or >1% hemoglobin reads. To reduce contamination from ambient myelin debris, cells with >3% myelin-associated reads were also excluded. Nuclei were retained if they expressed between 400 and 12,000 genes and had >700 unique molecular identifiers (UMIs). For doublet identification, scDblFinder was utilized to detect and remove artificial doublets[29]; clusters exhibiting high mitochondrial content (>95th percentile within the cluster) were further filtered to ensure dataset purity.

Normalization and variance stabilization were performed using SCTransform (v2), regressing out the percentage of mitochondrial reads. 3,000 highly variable genes were selected for downstream analysis. Dimensionality reduction was performed using t-distributed Stochastic Neighbor Embedding (tSNE) on the first 30 principal components. Graph-based clustering was performed at a resolution of 1.0. Integration of samples was achieved using Seurat’s SCT-based integration workflow to correct for batch effects and align shared biological states across conditions.

### Cell-Type Annotation

Mouse hippocampal cell types were annotated manually using established gene markers as listed in Figure 3B. The following markers were used to define major cell classes: Excitatory neurons: *Camk2a, Slc17a7, Tbr1*; Inhibitory neurons: *Gad1, Gad2, Sst, Vip, Pvalb*; Immature neurons: *Dcx, Neurod1, Tubb3*; Astrocytes: *Gfap, Aqp4, Aldh1l1, Slc1a3*; Oligodendrocytes: *Mog, Mbp, Plp1*; Oligodendrocyte precursor cells (OPCs): *Pdgfra, Cspg4, Sox10*; Microglia: *Cx3cr1, P2ry12, Tmem119, Aif1*; Endothelial cells: *Pecam1, Cldn5, Flt1*; Pericytes: *Pdgfrb, Rgs5, Vtn*; Fibroblasts: *Col1a1, Col1a2, Dcn*; Ependymal cells: *Foxj1, Tmem212*; Choroid plexus epithelial cells: *Ttr, Aqp1*. Excitatory neuron subtypes (e.g., CA1, CA3, DG granular) were further resolved based on cluster-specific enrichment and spatial identity inferred from cluster topology.

### Differential Gene Expression and Functional Analysis

Differentially expressed genes (DEGs) were identified using the FindMarkers function in Seurat via the Wilcoxon rank-sum test, as described previously [16, 22, 30]. Significant DEGs were defined by an adjusted p-value < 0.05 and an absolute log_2 fold-change > 0.5. Functional enrichment analyses, including Gene Ontology (GO), were performed using the clusterProfiler R package (version 4.10.1) and the org.Mm.eg.db database (version 3.18.0) with default parameters. Additionally, Gene Set Enrichment Analysis (GSEA) was conducted utilizing the Molecular Signatures Database (MSigDB) collection (curated gene sets). Enrichment scores were calculated within individual categories (e.g., KEGG, GO), and the results were integrated for comprehensive visualization.

### Trajectory Inference

To model astrocyte state transitions, trajectory analysis was performed using Monocle3. The Seurat object containing astrocyte clusters was converted to a subset. The trajectory graph was learned using learn_graph, and cells were ordered in pseudotime using order_cells, with the root node defined by the homeostatic cluster 0.

### Cell-Cell Communication Analysis

Intercellular communication networks were inferred using the CellChat R package (version 2.1). Interaction probabilities were computed based on the expression of known ligand-receptor pairs in the CellChatDB mouse database [31]. Significant interactions were identified using a permutation test, and communication strength was visualized using heatmaps to compare signaling pathways between WT and aKO conditions.

### Co-Expression Network Analysis (hdWGCNA)

To identify gene co-expression modules associated with specific cell states, high-dimensional Weighted Gene Co-expression Network Analysis (hdWGCNA) was applied. Metacells were constructed to reduce sparsity, and a soft power threshold was selected to achieve a scale-free topology. Co-expression modules were constructed, and module eigengenes were correlated with experimental conditions (Genotype and Treatment) to identify disease-associated modules.

### Gene Regulatory Network Analysis

To infer transcription factor (TF) activity from single-nucleus gene expression data, we utilized the VIPER (Virtual Inference of Protein-activity by Enriched Regulon analysis) method[32] implemented in the decoupleR R package. A comprehensive gene regulatory network (regulon) containing signed TF-target interactions was obtained from the CollecTRI resource (mouse organism) via OmnipathR[33]. CollecTRI integrates multiple resources of curated regulons to provide broad coverage of transcriptional interactions [34].

Using this prior knowledge network, the gene expression matrix (log-normalized counts) was transformed into a protein activity matrix. The run viper function estimates the activity of each TF based on the enrichment of its downstream target genes within the gene expression signature of each individual nucleus. The resulting Normalized Enrichment Scores (NES) were used to quantify relative TF activity, where positive scores indicate activation and negative scores indicate repression.

### Statistical Analysis

Statistical analyses were mainly performed using OriginPro software (OriginPro 2020). Data are presented as mean ± standard error (SEM). Two-group comparisons were analyzed using the One-way ANOVA or Mann-Whitney U test (for non-parametric data). Multiple group comparisons were analyzed using One-way or Two-way Analysis of Variance (ANOVA) followed by Tukey’s post-hoc test. Time-course data (Racine scores) were analyzed using Two-way repeated measures ANOVA. Survival curves (seizure incidence) were analyzed using the Kaplan-Meier method with the Log-rank (Mantel-Cox) test or Holm-Sidak multiple comparisons. Proportions were compared using the Chi-square test. A p-value < 0.05 was considered statistically significant.

## Results

### Astrocytic *mgll* deletion confers robust neuroprotection against kainic acid-induced status epilepticus

To determine the cellular source of MAGL-mediated neuroprotection, we compared wild-type (WT) mice with astrocyte-specific (aKO), neuron-specific (nKO), and total (tKO) *mgll* knockout mice to define the relative contributions of 2-AG signaling in astrocytes versus neurons to seizure activity (Fig. 1A). Upon administration with KA, both tKO and aKO mice exhibited marked resistance to seizure induction, characterized by a significantly prolonged latency to the first generalized seizure (Racine stages 4∼5) (Fig. 1B). Furthermore, the maximum Racine score and the percentage of seizure onset within one hour were significantly reduced in aKO and tKO mice compared to WT controls (Fig. 1C, S1A). In sharp contrast, neuron-specific deletion alone was insufficient to confer comparable resistance; nKO mice displayed seizure severity and latency profiles statistically similar to WT controls (Fig. 1B&C). These findings suggest that while neuronal MAGL may contribute to baseline synaptic buffering, the potent anti-seizure efficacy observed in the global knockout (tKO) is driven primarily by inactivation of *mgll* in astrocytes. To assess the functional consequences of SE, we performed open field testing. Consistent with KA-induced excitotoxicity, WT and nKO mice displayed marked hyperactivity, traveling significantly longer distances and less freezing time compared to vehicle controls. However, aKO and tKO mice exhibited locomotor profiles indistinguishable from baseline, indicating preservation of normal network function (Fig. 1D; Fig. S1B). We observed that while aKO effectively blocked seizure-induced neurodegeneration (Fig. 1E), the lack of protection in nKO mice suggests that the potent anti-seizure efficacy observed in tKO is driven primarily by astrocytic 2-AG signaling rather than neuronal signaling. Finally, to validate that the increased seizure susceptibility observed in nKO mice was driven by elevated 2-AG levels in neurons in the presence of intact astrocytic MAGL activity, and that the resilience of tKO mice was conferred by enhanced 2-AG signaling in astrocytes, we treated nKO mice with JZL-184, a potent and selective MAGL inhibitor[17]. Pharmacological inhibition of the remaining non-neuronal MAGL in nKO mice effectively rescued the phenotype, significantly prolonging the latency to seizure onset (Fig. 1F). These findings support the conclusion that the protective effects of global MAGL knockout against kainic acid-induced seizures arise primarily from inactivation of MAGL in astrocytes.

**Figure 1.**
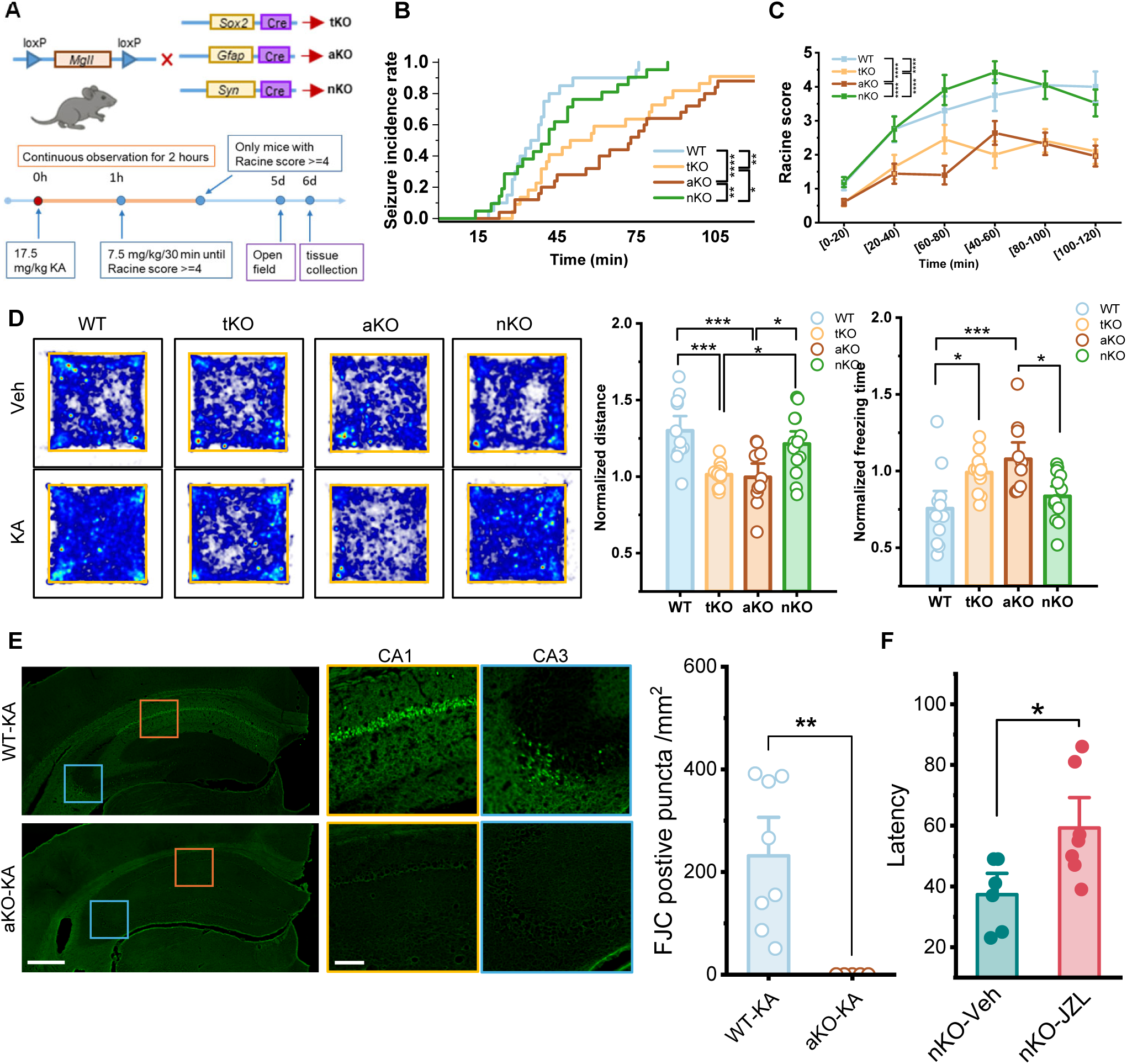
Astrocyte-specific *mgll* deletion attenuates kainic acid (KA)-induced seizure susceptibility and hippocampal neurodegeneration (A) Schematic representation of the genetic strategy used to generate total *mgll* knockout (tKO) mice (using Sox2-Cre), astrocyte-specific *mgll* knockout (aKO) mice (using Gfap-Cre), and neuron-specific *mgll* knockout (nKO) mice (using Syn-Cre). (B) Kaplan-Meier curves showing the cumulative seizure incidence rate following KA administration in WT, tKO, aKO, and nKO mice. Statistical significance was determined using the Kaplan-Meier estimator with Holm-Sidak’s multiple comparison test. (C) Time course of seizure severity quantified by the Racine scale over 2 hours post-KA injection. Data were analyzed using two-way repeated measures ANOVA followed by Tukey multiple comparison test. (D) Representative heatmaps illustrating the activity tracks of mice in the Open Field test. Warmer colors indicate strictly higher time spent in that location. Quantification of locomotor and anxiety-like behaviors in the Open Field test, showing normalized total distance traveled and normalized freezing time. Data are means ± SEM. *p <0.05, **p<0.01, ***p< 0.001 (n=11∼13/group, ANOVA with post hoc Turkey test). (E) Representative immunofluorescence images of Fluoro-Jade C (FJC) staining in the hippocampus of WT and aKO mice following KA treatment. High-magnification insets (colored boxes) display the CA1 and CA3 regions. Scale bars: 500 μm (overview) and 100 μm (insets). Quantification of FJC-positive puncta density (puncta/mm²) in the hippocampus, showing significantly reduced neurodegeneration in aKO mice compared to WT Data are means ± SEM. **P<0.01 (the Mann-Whitney test, n=4/group). (F) Latency to first seizure of nKO mice with JZL-184 or DMSO. Data are means ± SEM. *P<0.05 (n= 6-7/group, unpaired Student’s t-tes).

### Astrocytic MAGL serves as a critical regulator of KA-induced hippocampal neuroinflammation

To contextualize our findings within the broader biological landscape of epilepsy, we re-analyzed public bulk RNA-sequencing datasets from the hippocampal dentate gyrus (GSE213393)[35]. We observed distinct temporal dynamics in the endocannabinoid (eCB) system following KA-induced seizures (Fig. 2A). Interestingly, while *mgll* expression positively correlated with neuroinflammation scores in control and contralateral tissues, this relationship was reversed in the ipsilateral injured hippocampus (Fig. 2B). Several previous studies have shown that seizure activity enhances the production of 2-AG as a self-protective mechanism [36, 37]. This suggests that MAGL in the brain plays an important role as a compensatory adaptive response to acute neuroinflammation in maintaining brain homeostasis [38]. Further analysis of CA1 tissue (GSE99577)[39] showed a transient, biphasic response of *mgll* expression, which was downregulated first and then upregulated, along with a sustained increase of neuroinflammation score, particularly for the markers of astrocytes and microglia (Fig. S1D,E). This transient engagement suggests that the brain’s endogenous protective mechanism is insufficient to arrest the progression of gliosis.

**Figure 2.**
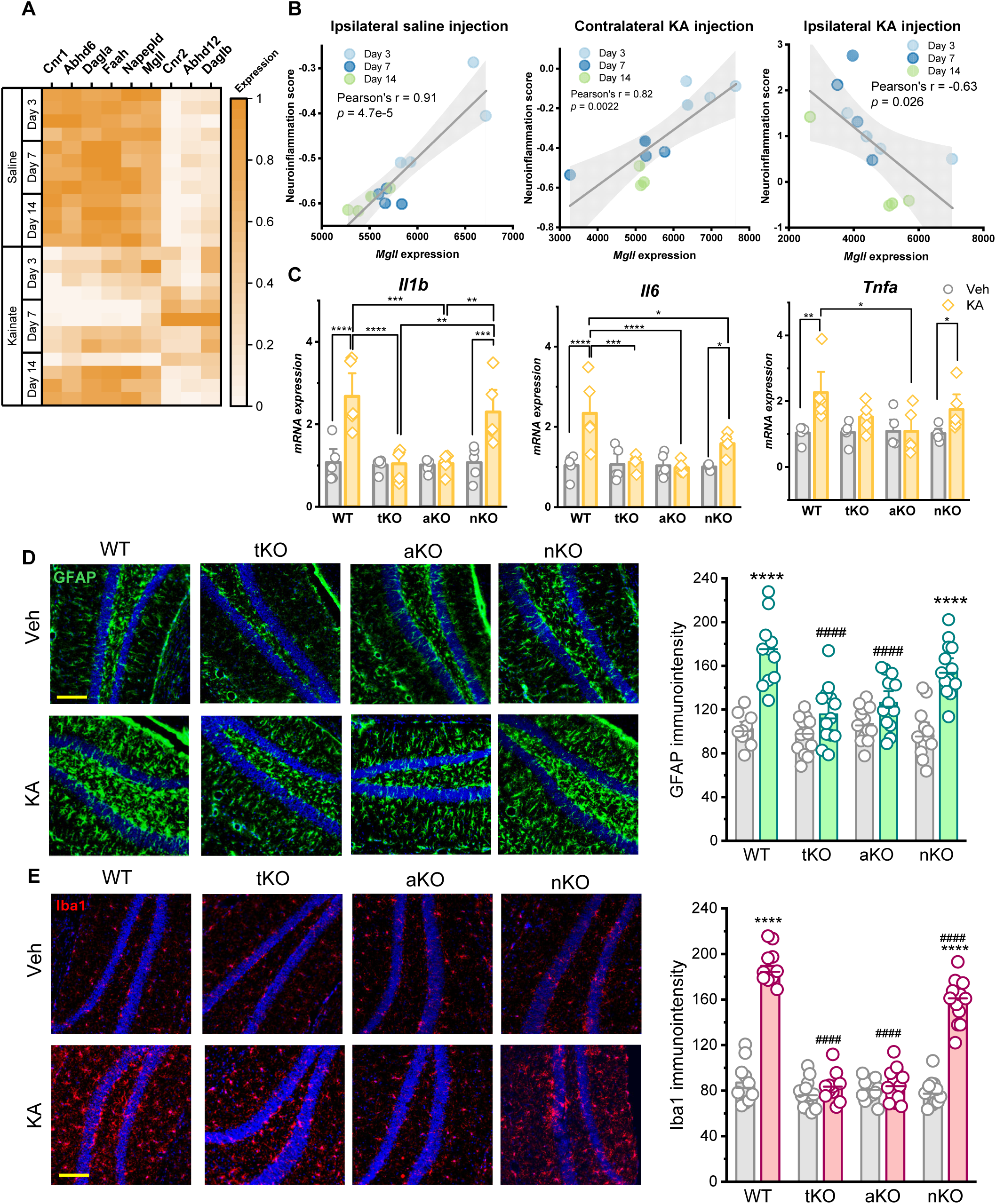
Comparison of seizure susceptibility, neuroinflammation, and behavioral deficits in global (tKO), astrocyte-specific (aKO), and neuron-specific (nKO) *mgll* knockout mice. (A) Heatmap illustrates the temporal mRNA expression profiles of the endocannabinoid system (eCB) in the hippocampus of mice treated with saline or KA at days 3, 7, and 14 post-injections. The color scale indicates relative expression levels. (B) Pearson correlation analysis between *mgll* expression levels and a composite neuroinflammation score (derived from the expression of *Aif1, Cx3cr1, Cd68, Il1b, Ccl2, Cxcl10, Ptgs2, Nos2, C1qa, C3, Itgam, and Serping1*) across different time points (Day 3, 7, 14) in ipsilateral saline-injected DG, contralateral KA-injected DG, and ipsilateral KA-injected DG. The linear regression significance was determined by ANOVA (p-values indicated on plots). (C) Relative mRNA expression levels of proinflammatory cytokines *Il1b*, *Il6*, and *Tnfa* in the hippocampus. Data are means ± SEM. *p<0.05, **p <0.01, ***p < 0.001, ****p < 0.001 (ANOVA with post hoc Turkey test, n=5/group). (D) Representative immunofluorescence images of GFAP staining (green) in the DG across different genotypes and treatments. Nuclei are stained with DAPI (blue). Scale bar: 100 μm. Quantification of GFAP immunofluorescence intensity in the DG regions (n=4-5 mice). Data are means ± SEM. *p<0.05, **p< 0.01, ****p<0.001, compared with Veh in each genotype; ^#^p<0.05, ^##^p<0.01, ^####^p<0.001 compared with WT-KA (ANOVA with Turkey test, 2∼3 sections/mice, n=4 mice/group)). (E) Representative immunofluorescence images of Iba1 staining (green) in the DG across different genotypes and treatments. Scale bar: 100 μm. Quantification of Iba1 immunofluorescence intensity in the DG regions (n=4-5 mice). Data represent individual slice values (2–3 slices per mouse) from n = 4 mice per group. Data are means ± SEM. *p<0.05, **p<0.01, ****p < 0.001 compared with Veh in each genotype, ^#^p<0.01, ^##^p<0.01, ^####^p<0.001 compared with WT-KA (ANOVA with Turkey test, n=4∼5 mice/group).

**Figure 3.**
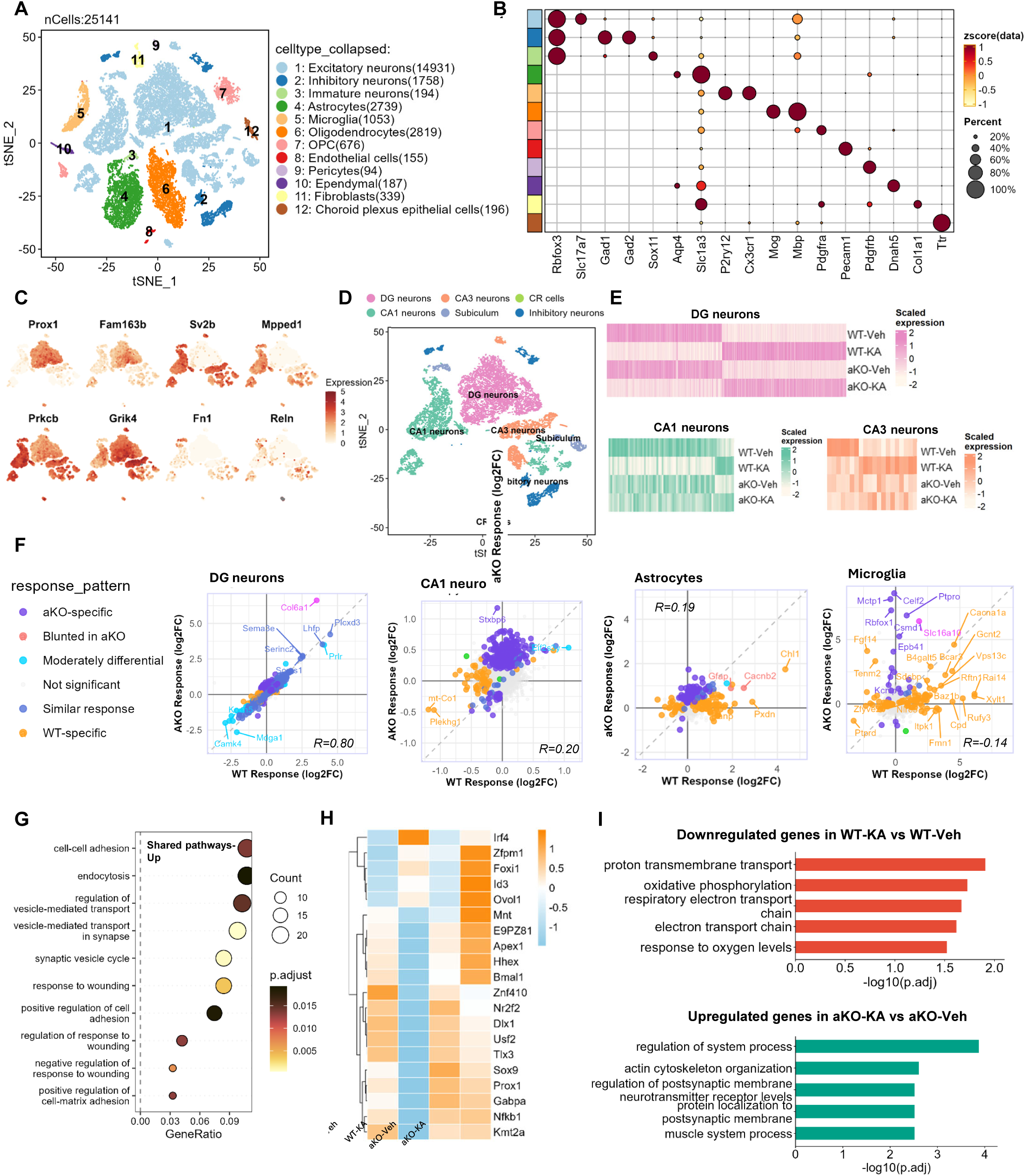
Single-nucleus RNA sequencing reveals distinct cell type-specific transcriptional responses to KA in aKO mice. (A) t-Distributed Stochastic Neighbor Embedding (t-SNE) visualization of nuclei isolated from the hippocampus, clustered into 12 distinct cell types. (B) Dot plot showing the expression of canonical marker genes used to identify major cell types. Dot size represents the percentage of cells expressing the gene, and color intensity represents the average expression level. (C) Feature plots displaying the expression of specific marker genes used to define neuronal subpopulations. (D) t-SNE plot showing the sub-clustering of neuronal populations into Dentate Gyrus (DG) granular cells, CA1 pyramidal neurons, CA3 pyramidal neurons, Subiculum, Inhibitory neurons, and Cajal-Retzius cells. (E) Heatmaps showing the relative expression of differentially expressed genes (DEGs) across experimental groups (WT-Veh, WT-KA, aKO-Veh, aKO-KA) in DG granular cells, CA1 pyramidal neurons, CA3 pyramidal neurons. (F) Scatter plots comparing the transcriptional response to KA (log2 Fold Change) between WT (x-axis) and aKO (y-axis) mice in DG granular cells, CA1 pyramidal neurons, astrocytes, and microglia. Points are colored based on their response pattern. Pearson correlation coefficients (R) are indicated for each cell type. (G) Gene Ontology (GO) enrichment analysis of pathways significantly upregulated in DG in both WT and aKO mice (Shared pathways Up) following KA treatment. Dot size indicates gene counts; Pathways represent adjusted p-value.(H) Heatmap displaying the expression of key transcription factors across the four experimental groups. (I) Bar charts showing significant GO terms for pathways downregulated in WT-KA mice and pathways upregulated in aKO-KA mice. The x-axis represents -log10 (adjusted p-value).

Thus, we examined whether this astrocyte-specific protection extended to the neuroinflammatory sequelae of SE. qPCR analysis demonstrated robust induction of proinflammatory cytokines, specifically *Il1b*, *Il6*, and *Tnfa*, in the hippocampi of WT and nKO mice (Fig. 2C). In contrast, this cytokine storm was significantly attenuated in aKO and tKO mice. Immunohistological analysis at Day 6 revealed that nKO mice, similar to WT controls, exhibited profound reactive gliosis, characterized by widespread upregulation of GFAP and Iba1. Astrogliosis is pervasive throughout the hippocampus, including DG, CA1 and CA3. In contrast, aKO and tKO mice maintained glial markers at near-basal levels (Fig. 2D, S2A&B). Iba1 was used to detect microgliosis. Activated microglia, characterized by hypertrophic cell bodies and thickened processes, were recruited to DG granular cell layer, showing an increased immunoinstensity in WT and nKO mice (Fig. 2E). The results of CA1 and CA3 are similar (Fig. S2C&D). These histological observations were confirmed by transcriptional profiling; Collectively, these data identify astrocytic 2-AG signaling as the dominant cellular mediators of MAGL-dependent protection against excitotoxic injury and inflammation.

### Astrocytic *mgll* deletion confers region- and cell type-specific resilience by preserving metabolic homeostasis and neuronal Identity

To dissect the molecular mechanisms and cellular heterogeneity underlying aKO-mediated neuroprotection, we performed single-nucleus RNA sequencing (snRNA-seq) on pooled bilateral hippocampal tissue from five mice per group (WT-Veh, WT-KA, aKO-Veh, aKO-KA). Only mice with Racine score reaching 4 and above after KA injection were selected. Following rigorous quality control, 25,141 high-quality nuclei were retained (Fig. 3A). Quality assessment are provided in Figures S3A&B. Unsupervised clustering and manifold learning resolved the cells into 12 major cell types: *Slc17a7* (excitatory neurons), *Gad1/Gad2* (inhibitory neurons), *Sox11*(immature neurons), *Slc1a3* (astrocytes), *P2ry12/Cxc3r1* (microglia), oligodendrocytes, oligodendrocyte precursor cells (OPC), endothelial cells, pericytes, ependymal cells, fibroblasts and choroid plexus epithelial cells (Fig. 3A). Canonical marker expression confirmed distinct molecular identities for all clusters (Fig. 3B). To resolve the diversity of excitatory neurons, we further identified Dentate Gyrus (DG) granule cells (*Prox1^+^, Fam163b*^+^), CA1 (*Sv2b*^+^, *Mpped1*^+^, *Prkcb*^+^)and CA3 (*Sv2b*^+^, *Grik4*^+^) pyramidal cells, Subiculum neurons (*Fn1*^+^), and Cajal-Retzius cells (*Reln*^+^) (Fig. 3C&D, Fig S3C). Given their central role in hippocampal excitability, we prioritized the analysis of principal excitatory types.

We first assessed the global impact of KA on transcriptional states. Interestingly, the response to KA was cell-type dependent. In DG granule cells, the response to seizure activity appeared highly conserved. Heatmap analysis revealed a striking concordance in expression profiles, with WT and aKO mice displaying nearly identical patterns of up- and downregulated gene clusters (Fig. 3E). This similarity was statistically quantified by scatter plot analysis of Log2FC values, where genes aligned tightly along the diagonal, yielding a high Pearson correlation coefficient (Pearson’s R =0.80) (Fig. 3F). In sharp contrast, the response in CA1/CA3 pyramidal cells was markedly divergent. In the vulnerable CA1 and CA3 pyramidal populations, aKO mice exhibited a blunted expression of WT-KA induced DEGs (Fig. 3E&F; Fig. S3D). This suggests that aKO confers a capacity for active transcriptional remodeling in pyramidal cells that is otherwise compromised in the wild-type condition. In inhibitory neurons, aKO showed resilience to KA-induced gene upregulation (Fig. S3E&F). This pattern of genotype-dependent resistance extended to the glial compartment; for example, the reactive astrocyte marker *Gfap* was significantly attenuated in aKO mice, and microglia also displayed a dynamic and distinct trajectory shift. This global data suggested that aKO does not simply suppress all signaling but specifically protects vulnerable populations (CA1) while preserving the baseline response in resistant populations (DG).

Finally, we assessed the subfield-specific neuronal responses. In DG granular cells, we identified a core set of pathways shared between aKO-KA and WT-KA (vs. their respective vehicles). These shared signatures confirmed that both genotypes responded to the excitatory impairment, mounting a common program of endocytosis and regulation of cell matrix (Fig. 3G). While global gene expression analysis suggested a conserved response to KA in the DG, regulatory network analysis based on transcriptional factors revealed a critical divergence in cellular resilience (Fig. 3H). In WT-KA mice, we observed a collapse of key granule cell identity and maintenance factors. Specifically, *Prox1*, the master regulator of DG neuronal identity, was markedly downregulated in WT-KA neurons but preserved in aKO-KA mice. Furthermore, aKO-KA mice exhibited sustained activity of neurogenic, and stem-cell associated factors (*Sox9*, *Id3*, *Nanog)* and metabolic regulators (*Tfam*, *Bmal1*) that were otherwise lost in the WT condition. Conversely, the inflammatory driver *Irf4* was activated in WT-KA neurons but blunted in the aKO. These data indicate that while aKO DG neurons sense the seizure stimulus, they are protected from the subsequent loss of cellular identity and metabolic failure.CA1 is a region classically vulnerable to metabolic and excitotoxic stress. In WT mice, KA triggered a bioenergetic crisis in CA1 neurons, defined by the extensive downregulation of oxidative phosphorylation and electron transport chain genes (Fig. 3I). In contrast, aKO mice exhibited a distinct, pro-survival response to KA. CA1 neurons in aKO mice exhibited a robust upregulation of pathways associated with structural plasticity, such as actin cytoskeleton organization (Fig. 3I). Analysis of upregulated pathways in WT-KA and downregulated pathways in aKO-KA revealed no significant GO enrichment (p.adjust > 0.05, Table S2), further suggesting that neurons in aKO mice maintain the capacity to remodel synaptic connections and reinforce synapse integrity in response to the seizure insult.

### Astrocytic *mgll* deletion alters astrocyte functional states and suppresses disease-associated microglia (DAM) activation

We then focused our analysis on astrocytes. Following additional quality control aimed at removing oligodendrocyte and microglial contamination, 2,671 astrocytes were retained and re-clustered into five transcriptionally distinct six subclusters (Fig. 4A). Notably, aKO mice contained a higher number of astrocytes than WT mice, in both Veh- and KA-treated conditions (Fig. 4B&C). Upon KA administration, WT mice exhibited a marked shift in astrocyte composition, characterized by the emergence of a KA-induced Cluster 5 and the reduce of Cluster 0. In contrast, aKO mice primarily expanded Cluster 2 (Fig. 4B&C). To resolve the heterogeneity of the astrocyte population, we first assessed the expression of canonical marker genes across the identified subclusters (Fig. 4D). It revealed distinct molecular identities corresponding to different functional states. Cluster 0 and Cluster 1 displayed high expression of homeostatic markers (*Slc1a2*, *Glul*, *Ndrg2*), suggesting a baseline or quiescent phenotype. In contrast, Cluster 5 exhibited a strong reactive and inflammatory profile, defined by the upregulation of *Gfap*, *Vim*, and immune-associated genes (*Stat3*, *Il1r1*). Cluster 4 was clearly identifiable as a proliferative population, exclusively expressing cell cycle markers such as *Mki67*, *Top2a*, and progenitor markers (*Prox1*, *Fabp7*). The Gene Set Enrichment Analysis (GSEA) corroborated our marker-based classification while revealing a deeper metabolic divergence between the subsets (Fig. S4A). Cluster 2 presented a unique bioenergetic profile. Unlike the glycolytic Cluster 1, Cluster 2 showed robust enrichment for oxidative phosphorylation and ATP biosynthetic processes, alongside immunomodulatory pathways like interleukins 4 and 13 signaling. The proliferative identity of Cluster 4 was functionally confirmed by the massive upregulation of cell cycle. Cluster 5, aligning with its inflammatory marker profile, showed enrichment for cellular response to hypoxia and axon guidance, suggesting these astrocytes are responding to acute tissue injury and environmental stress.

**Figure 4.**
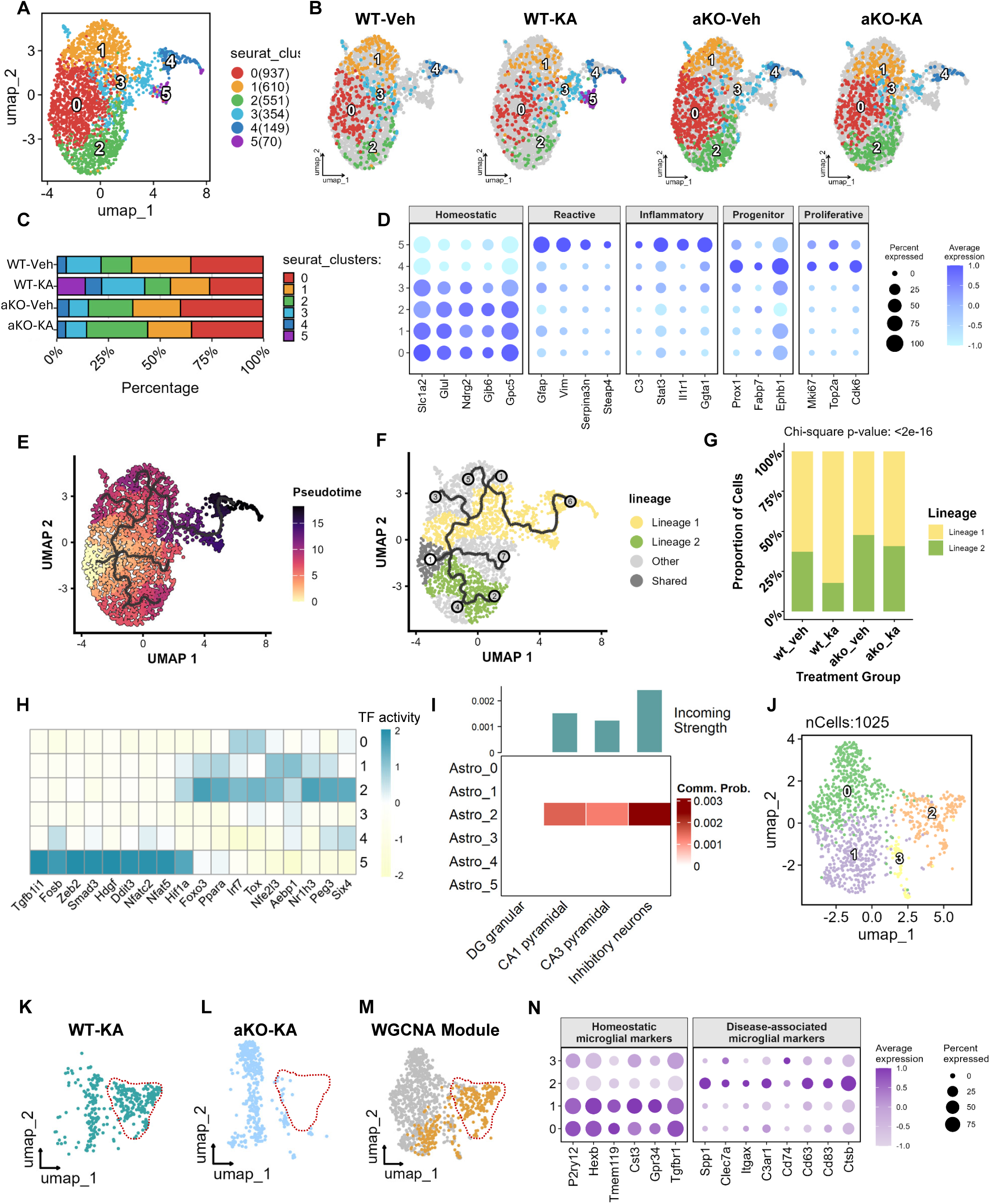
*Mgll* deletion alters astrocyte functional states and suppresses disease-associated microglia (DAM) activation. (A) UMAP visualization of astrocytes clustered into 6 distinct subclusters (Clusters 0∼5). (B) UMAP plots split by experimental group (WT-Veh, WT-KA, aKO-Veh, and aKO-KA), visualizing the distribution of astrocyte subclusters across conditions. (C) Stacked bar chart showing the percentage composition of each astrocyte subcluster within the four experimental groups. (D) Dot plot of canonical marker genes used to annotate astrocyte states: Homeostatic, Reactive, Inflammatory, Progenitor, and Proliferative. Dot size represents the percentage of expressing cells, and color intensity indicates average expression levels. (E) Pseudotime trajectory analysis of astrocyte subpopulations, colored by pseudotime values. (F) UMAP showing the bifurcation of astrocyte trajectories into two distinct lineages (Lineage 1 in yellow and Lineage 2 in green). (G) Quantification of the proportion of cells belonging to Lineage 1 versus Lineage 2 across experimental groups. Statistical significance was determined using the Chi-square test. (H) Heatmap displaying the activity of key transcription factor regulons across astrocyte clusters (0∼5). (I) Cell-cell communication analysis (CellChat) illustrating the communication probability between astrocyte subclusters (rows) and neuronal populations (columns). (J) UMAP visualization of microglia clustered into distinct subpopulations. (K∼M) UMAP plots highlight a specific disease-associated microglial module identified by hdWGCNA. The red dashed line indicates a population of activated microglia present in WT-KA (K) but largely absent in aKO-KA (L), corresponding to the hdWGCNA module (M). (N) Dot plot showing the expression of Homeostatic microglial markers and Disease-associated microglial (DAM) markers across microglial clusters.

To determine the dynamic trajectories underlying these responses, we performed pseudotime analysis (Fig. 4E). Astrocytes displayed multipotent reactivity (Fig. 4F). Both WT and aKO switched into lineage 1, where switched into inflammatory Cluster 5 (Fig. S4B), while aKO astrocytes preferentially accumulated in lineage 2, leading to Cluster 2 (Fig. 4F&G). Density plot along pseudotime suggested that aKO astrocytes positioned within Cluster 2 (Fig. S4C). To understand this bifurcation, direct comparison of transcriptional factors between Cluster 2 and Cluster 5 demonstrated that aKO cells activates a completely different set of regulators focused on survival (e.g., *Foxo3, Tox*) and lipid metabolism (e.g., *Ppara*, *Aebp1*), suggesting a retained capacity to support neuronal homeostasis (Fig 4H). We further inferred intercellular communication networks using CellChat. We specifically examined the endocannabinoid signaling pathway (2-AG–Cnr1). Remarkably, among all astrocyte subpopulations, only Cluster 2 exhibited significant communication probabilities with neuronal targets (Fig. 4I), which can be contributed to the enrichment of *Dagla* (diacylglycerol lipase-α, the synthetase of 2-AG) expressing astrocytes (Fig. S4D).

Given the intimate functional coupling between astrocytes and microglia in neuroinflammation, we next interrogated the microglial landscape. Re-clustering of microglia yielded four distinct subclusters (Fig. 4J). In WT mice, KA injection triggered a profound emergence of a specific disease-associated population, Cluster 2, a shift that was significantly attenuated in aKO mice (Fig. 4K&L, S5A&B). WGCNA identified a specific gene module predominantly expressed in Cluster 2 (Fig. 4M, S5C), which was significantly enriched for pathways involving apoptosis and cytokine signaling (Fig. S5D). The result is confirmed by canonical markers. Cluster 0, 1, and 3 maintained canonical surveillance markers (e.g., *P2ry12*, *Cx3cr1*), consistent with a homeostatic phenotype. In contrast, the WT-dominant Cluster 2 displayed a pro-inflammatory signature characterized by high expression of *Spp1* and *Cd63*, a profile aligning with Disease-Associated Microglia (DAM) previously implicated in aberrant synaptic pruning (Fig. 4N). Pseudobulk analysis confirmed this genotype-specific divergence: aKO-KA microglia exhibited a blunted induction of the DAM signature while sustaining high expression of the homeostatic program, suggesting that aKO prevents the polarization of microglia into an inflammatory state (Fig. S5E).

### MAGL inhibition attenuates seizure susceptibility and hippocampal neuroinflammation are mediated via CB1-PPAR-**γ** signaling pathways

To determine the signaling pathways downstream of MAGL inhibition, we first investigated the requirement of the CB1 receptor. We administered the specific MAGL inhibitor JZL-184 to *Cnr1* knockout (Cnr1^−/-^) mice prior to KA induction (Fig. S6A, B). In the absence of CB1 receptor, the neuroprotective effects of JZL-184 were completely abolished; we observed no significant extension of seizure latency (Fig. 5A&B) and no reduction in seizure severity (Fig. 5C) compared to vehicle-treated controls. Furthermore, JZL-184 failed to attenuate reactive astrogliosis (GFAP) or microglial activation (Iba1) in the DG (Fig. 5D&F) or the CA1 and CA3 subfields (Fig. S6C&D). This widespread lack of rescue confirms that CB1 signaling is a prerequisite for 2-AG-mediated neuroprotection. Transcriptomic profiling indicated robust metabolic reprogramming in aKO astrocytes is represented by PPAR signaling pathways. Although our analysis specifically highlighted PPARα activity, the endocannabinoid 2-AG has been demonstrated to resolve neuroinflammation by activating PPAR-γ [16, 19, 40]. To determine if this specific pathway drives the protective phenotype, we treated wild-type mice with JZL-184 in the presence of the selective PPAR-γ antagonist GW9662 (Fig. 5G). The concurrent blockade of PPAR-γ effectively reversed the beneficial effects of JZL-184 on seizure severity, driving Racine scores back to vehicle-control levels (Fig. 5H&I). Similarly, GW9662 treatment negated the anti-inflammatory capability of JZL-184, restoring high levels of GFAP and Iba1 expression in the hippocampus (Fig. 5J& K). This suggested that the CB1-PPAR-γ signaling pathway is a primary mediator of seizure attenuation and suppression of neuroinflammation.

**Figure 5.**
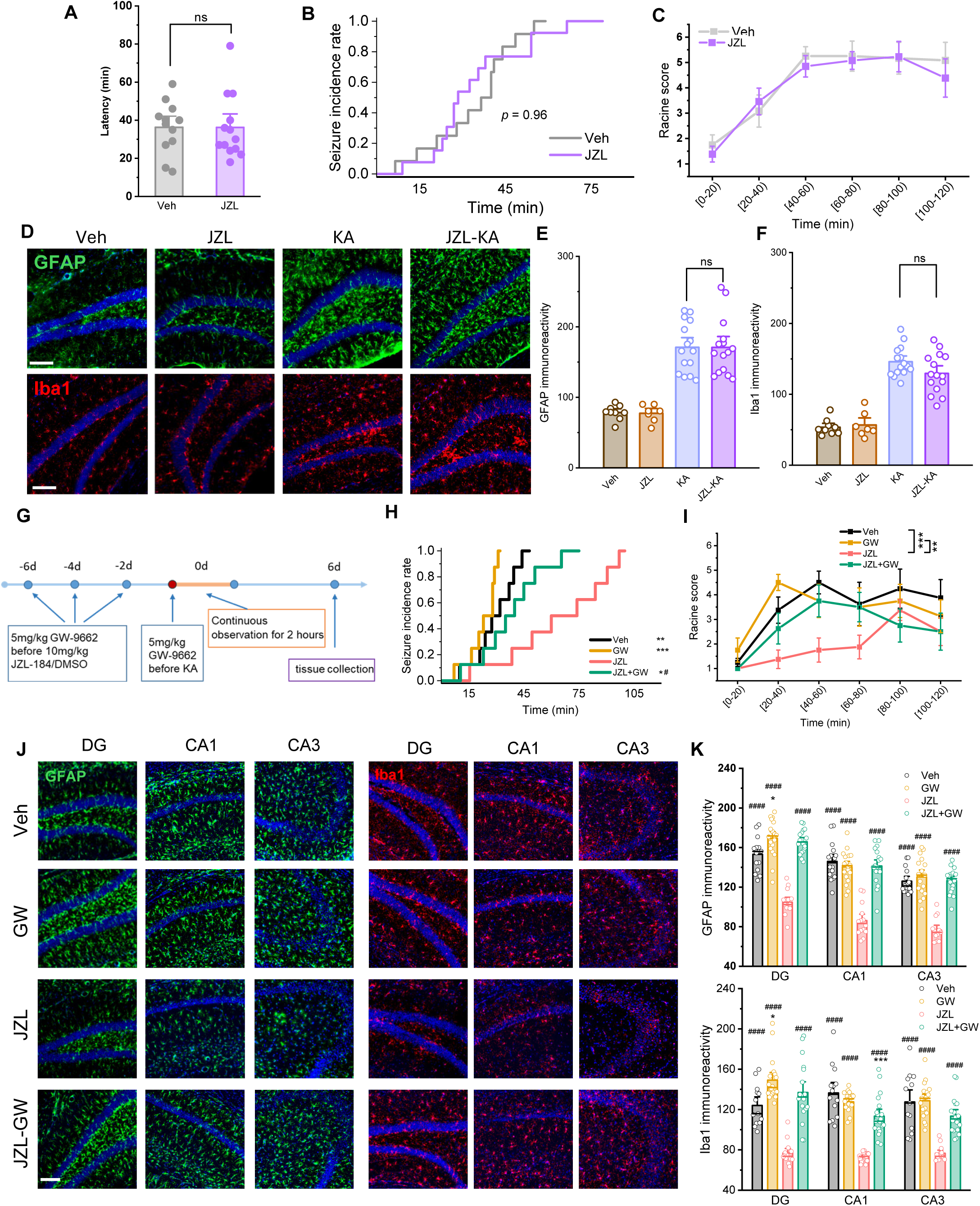
The neuroprotective effects of MAGL inhibition are mediated via CB1 receptors and PPAR-γ signaling pathways. (A) Assessment of seizure susceptibility in CB1 receptor knockout mice treated with the MAGL inhibitor JZL-184 compared to vehicle controls. Data are means ± SEM. (n=12∼13 mice/group). (B) Cumulative seizure incidence rate (Log-rank test). (C) Time course of Racine scores (Two-way repeated measures ANOVA). (D) Representative immunofluorescence images of GFAP (green) and Iba1 (red) staining in the hippocampus of DMSO- and JZL-treated mice following KA or saline (Veh) administration. Scale bar: 100 μm. (E&F) Quantification of GFAP (E) and Iba1 (F) immunoreactivity. Data are means ± SEM. (ANOVA with Turkey test, 3∼4 sections/animal, n=3∼4 mice/group). (G) Schematic timeline of the rescue experiment using GW-9662 (GW). (H) Kaplan-Meier curves showing cumulative seizure incidence (n=8). Statistical significance was determined by the Log-rank test with Holm-Sidak’s multiple comparison test. (I) Time course of seizure severity (Racine score). **p<0.01, ***p<0.001 (two-way repeated measures ANOVA with Turkey test, n=8 animals/group). (J) Representative immunofluorescence images of GFAP (green) and Iba1 (red) in the DG, CA1, and CA3 regions across the four treatment groups (Veh, GW, JZL, and JZL+GW). Scale bar: 100 μm. (K) Quantification of GFAP (top) and Iba1 (bottom) immunoreactivity in the hippocampal subregions. Data represent individual slice values (3–4 slices per mouse) from n = 4-5 mice per group. Data are means ± SEM. *p<0.05, **p<0.01, ****p<0.001 compared with Veh, ^#^p<0.05, ^##^p<0.01, ^####^p<0.001 compared with JZL (ANOVA with post hoc Turkey test, 3∼4 sections/animals, n=4∼5 animals/group).

### Astrocytic *mgll* deletion confers sustained protection against chronic behavioral deficits and neuronal loss

To evaluate whether the neuroprotective effects of astrocytic *mgll* deletion translate into long-term functional recovery, we assessed behavioral and histological outcomes in the chronic phase (Day 21). We first examined locomotor activity and anxiety-like behavior using the Open Field Test. WT-KA mice exhibited significant hyperactivity, characterized by increased total distance moved and mean activity, alongside a marked reduction in freezing time compared to vehicle controls (Fig. 6A&B). Strikingly, this behavioral phenotype was completely abrogated in aKO-KA mice, which displayed locomotor profiles indistinguishable from baseline vehicle controls. We next assessed cognitive function using the Novel Object Recognition (NOR) task. While WT-KA mice displayed a severe deficit in recognition memory, indicated by a significantly reduced recognition index, aKO-KA mice retained intact cognitive performance (Fig. 6C), suggesting the successful preservation of hippocampal memory circuits. Consistent with our snRNA-seq findings linking aKO astrocytes to synaptic assembly pathways, we evaluated both neuronal survival and synaptic integrity. Immunofluorescence staining for NeuN revealed that WT-KA mice suffered extensive neuronal loss in the CA1 region, whereas aKO-KA mice displayed robust preservation of neuronal density (Fig. 6D). Furthermore, we quantified the density of the postsynaptic marker PSD-95. WT-KA mice suffered a profound loss of synaptic puncta in the hippocampus; however, this synaptic erosion was effectively prevented in aKO-KA mice, which maintained PSD-95 density at levels comparable to vehicle controls. (Fig. 6E).

**Figure 6.**
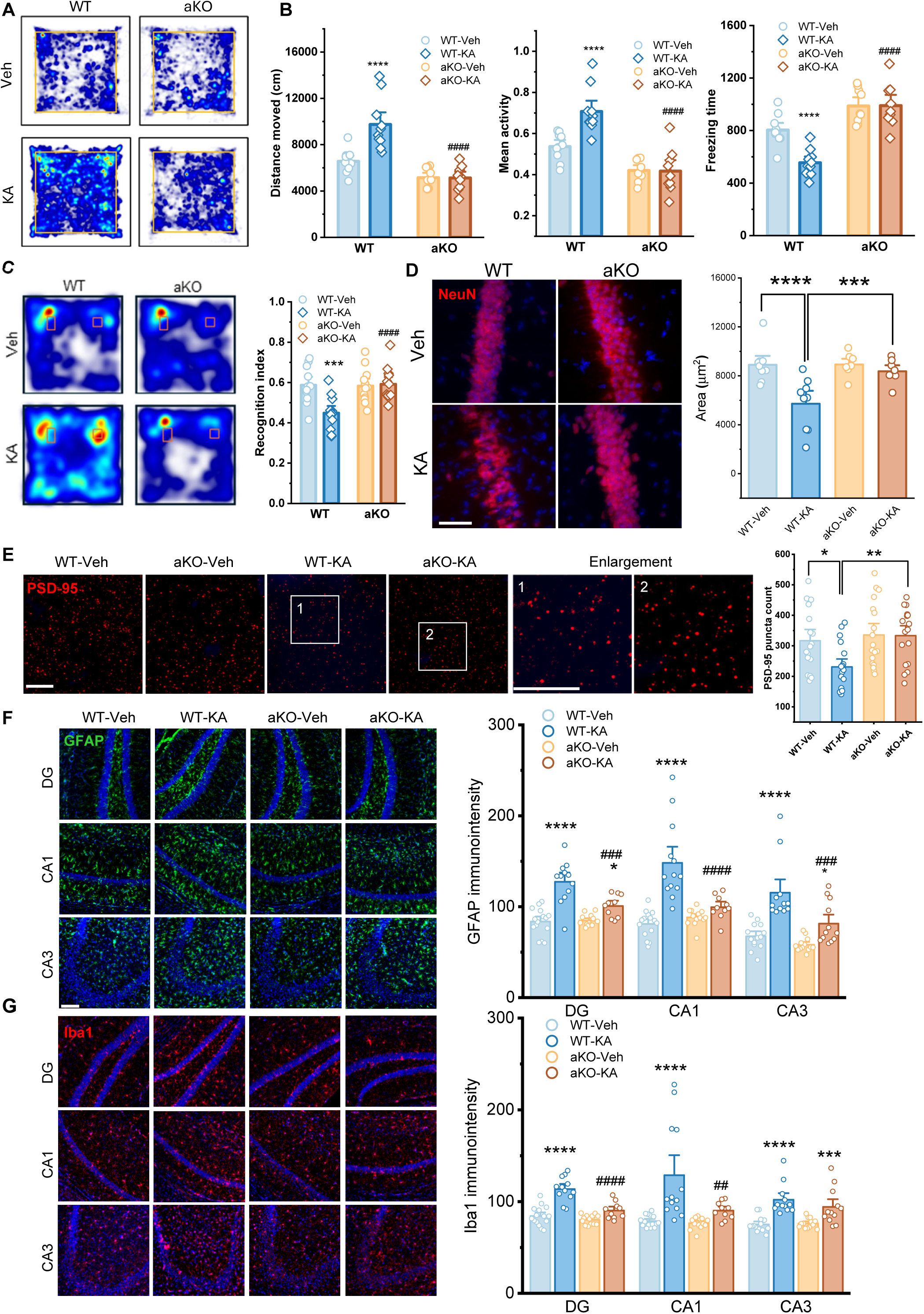
Astrocyte-specific *mgll* deletion prevents KA-induced cognitive deficits and synaptic loss. (A) Representative heatmaps illustrating the activity tracks of mice in the Open Field test. Warmer colors indicate strictly more time spent in that location. (B) Assessment of hyperactive activity and anxiety-like behavior in the Open Field test. Quantification of total distance moved, mean activity, and freezing time in WT and aKO mice treated with Vehicle or KA. Data are mean ± SEM. ****p<0.001 compared with Veh, ^####^p<0.001 compared with WT-KA (ANOVA with Turkey test, n=9∼10 animals/group). (C) Representative heatmaps of the Novel Object Recognition (NOR) test, illustrating the exploration time spent around the objects. Quantification of the Recognition Index is shown as mean ± SEM. ***p < 0.001 compared with Veh in each genotype, ####p < 0.001 compared with WT-KA (ANOVA with post hoc Turkey test, n=11∼12 animals/group). (D) Representative images of NeuN (red) staining in the CA1 region with quantification. Scale bars: 50 μm. Data are means ± SEM. ***p< 0.001, ****p<0.0001 (ANOVA with post hoc Turkey test, 2∼3 images/animals, n=3 animas/group). (E) Representative immunofluorescence images of the postsynaptic marker PSD-95 (red) in the hippocampus (CA1 region). Numbered boxes (1, 2) correspond to the enlargement insets on the right. Scale bars: 10 μm. Data points in the quantification of PSD-95 positive puncta represent individual imaging fields. Data are means ± SEM. *p<0.05 (ANOVA with post hoc Turkey test, 4∼5 image/animals, n=4 animals/group). (F-G) Representative immunofluorescence images of GFAP (green, H) and Iba1 (red, I) staining in the DG, CA1, and CA3 regions of the hippocampus with quantification. Scale bar: 100 μm. Data are means ± SEM. *p<0.05, ****p<0.0001 compared with Veh; ^##^p<0.01, ^###^p<0.001, ^####^p < 0.0001 compared with WT-KA (ANOVA with post hoc Turkey test, 3∼4 image/animals, n=4 animals/group).

To determine if this functional rescue corresponded to a suppression of chronic pathology, we evaluated neuroinflammation and synaptic integrity. Immunofluorescence analysis revealed that WT-KA mice sustained widespread reactive astrogliosis and microglial activation throughout the DG, CA1, and CA3 subfields at Day 21. In contrast, aKO-KA mice exhibited significantly attenuated glial activation across all hippocampal regions (Fig. 6F&G), confirming that the early engagement of the astrocytic protective axis prevents the establishment of chronic neuroinflammation. Collectively, these data demonstrate that the astrocyte-specific modulation of 2-AG signaling provides durable neuroprotection, preserving synaptic structure and cognitive function long after the initial insult.

## Discussion

In the present study, we identify astrocytic MAGL as a critical metabolic gatekeeper that regulates the transition from adaptive neural responses to catastrophic excitotoxicity and chronic neuroinflammation associated with seizure activity. By combining cell type-specific genetic knockouts with single-nucleus transcriptomics, we demonstrate that the neuroprotective efficacy of MAGL inactivation is driven primarily by astrocytic 2-AG signaling rather than neuronal 2-AG signaling. This protection is not merely a consequence of reduced seizure burden but appears to stem from a fundamental reprogramming of the glial inflammatory response. While previous studies have established that systemic MAGL inhibition suppresses seizures through enhanced endocannabinoid (eCB) signaling, it has generally been assumed that this protection arises from direct excitatory neuron communication [7, 13], which can thereby suppress glutamatergic transmission at the synapse [41]. Our data challenges this neuron-centric view. We observed that neuron-specific *mgll* deletion failed to replicate the protective phenotype of global or astrocyte-specific *mgll* knockout. Neurons express high levels of presynaptic MAGL that terminates retrograde endocannabinoid signaling and modulates synaptic transmission and plasticity. However, recent evidence indicates that neuron-derived 2-AG selectively signals to presynaptic CB1R within a restricted synaptic microdomain, rather than diffusing to activate astrocytic CB1R [42]. While neurons are the primary source of synaptic 2-AG, astrocytic MAGL is principally responsible for generating 2-AG–derived arachidonic acid (AA) to regulate inflammatory processes [43]. Consistent with this spatial compartmentalization, prior studies have demonstrated that deletion of neuronal MAGL does not alter seizure susceptibility [44]. This dissociation suggests that the therapeutic potential of 2-AG mobilization lies not merely in dampening synaptic release, but in its capacity to resolve the profound neuroinflammatory response that accompanies status epilepticus (SE). These findings parallel recent observations in traumatic brain injury, where astrocytic, but not neuronal, 2-AG signaling was found to be critical for reducing secondary injury cascades [16].

Specifically, as SE typically triggers a rapid induction of proinflammatory cytokines, lowering seizure thresholds and promoting secondary post-seizure injury [2, 45], we show that astrocytic *mgll* deletion (aKO) averts the cytokine storm and reactive gliosis typically triggered by SE, thereby preserving hippocampal structural integrity, synaptic density, and long-term cognitive function. In our study, while wild-type (WT) and neuron-specific knockout (nKO) mice exhibited severe upregulation of these cytokines and extensive reactive gliosis, aKO mice maintained an inflammatory profile comparable to baseline. These results align with emerging evidence that astrocytes act as the primary immune sensors mediated by 2-AG signaling in the epileptic brain, capable of orchestrating both the initiation and resolution of neuroinflammatory responses through their extensive spatial coverage and metabolic coupling with neurons [46–49].

The significant phenotypic differences observed between nKO and aKO mice strongly imply that astrocytic 2-AG acts as an important endogenous mediator that terminates neuroinflammatory responses to excitotoxic insults. By deleting *Mgll* in astrocytes, we likely increase the local availability of 2-AG, which engages a feed-forward protective mechanism that arrests the polarization of astrocytes into neurotoxic states. This is consistent with previous reports indicating that eCB signaling can suppress astrocyte activation in models of traumatic encephalopathy[23] and Alzheimer’s disease[50], extending this protective role to the acute excitotoxic environment of epilepsy. Moreover, the spatial compartmentalization of 2-AG signaling may be particularly important in this context. Unlike systemically administered cannabinoids that activate CB1 receptors throughout the brain, astrocytic MAGL deletion may create microdomains of elevated 2-AG specifically in perisynaptic astrocytic processes, allowing for precise spatiotemporal control of both neuronal excitability and glial reactivity[43].

Our single-nucleus RNA sequencing (snRNA-seq) analysis provides a high-resolution mechanistic explanation for this anti-inflammatory phenotype. In WT mice, KA induction drove astrocytes toward a reactive and inflammatory trajectory (Cluster 5) and microglia toward a Disease-Associated Microglia (DAM) phenotype (Cluster 2) implicated in aberrant synaptic pruning[51]. Strikingly, aKO astrocytes largely bypassed this inflammatory fate, instead populating a distinct metabolic state (Cluster 2) characterized by enhanced oxidative phosphorylation and lipid metabolism pathways. This metabolic reprogramming resembles the neuroprotective astrocyte phenotype, where astrocytes upregulate bioenergetic support to sustain neurons during stress[52, 53]. This suggests that *Mgll* deletion does not simply silence astrocytes but actively repurposes them to support neuronal bioenergetics during stress, potentially through enhanced lactate shuttle activity and provision of metabolic substrates[54]. Furthermore, the blunted DAM signature in microglia of aKO mice indicates that non-cell-autonomous signaling from protected astrocytes is related to limited microglial activation. This finding is consistent with the emerging concept that astrocyte-microglia crosstalk determines neuroinflammation in various diseases[55, 56].

Mechanistically, our pharmacological rescue experiments clarify the signaling hierarchy underlying this protection. The complete loss of efficacy in CB1 knockout mice confirms that CB1 receptor activation is the prerequisite for 2-AG-mediated protection, consistent with extensive literature demonstrating CB1 as the primary mediator of endocannabinoid effects in the brain. However, the partial divergence observed with PPAR-γ antagonism offers a nuanced view of the intracellular signaling cascades. While PPAR-γ blockade fully reversed the anti-inflammatory benefits and increased seizure severity, it did not completely abolish the delay in seizure onset. This finding supports a hypothesis of dual-mechanism of endocannabinoid signaling: an immediate, membrane-mediated modulation of excitability, followed by a sustained, anti-inflammatory genomic effect mediated by the intracellular CB1-PPAR-γ axis. The rapid, membrane-proximal effects likely involve CB1-mediated inhibition of adenylyl cyclase, reduction of cAMP levels, and suppression of presynaptic calcium influx[57, 58], while the slower PPAR-γ-dependent effects involve transcriptional reprogramming[59]. It is this secondary axis that appears crucial for preventing the cascade of reactive gliosis and secondary neurodegeneration. Previous studies also showed that the anti-neuroinflammation effect of astrocyte-specific MGL-deficiency cannot be fully abrogated by the inverse CB1receptor agonist[60]. By engaging nuclear receptors, accumulated endocannabinoids likely repress NFκB and other inflammatory transcription factors[18, 61], preventing the transformation of the hippocampus into a pro-inflammatory environment that facilitates secondary neurodegeneration. Indeed, PPAR-γ activation has been shown to suppress the expression of COX-2, iNOS, and pro-inflammatory cytokines in epilepsy, providing a plausible molecular mechanism for the anti-inflammatory phenotype we observed[62].

The downstream consequences of this immunometabolic rescue are evident in the preservation of neuronal viability and identity. The CA1 subfield, typically vulnerable to metabolic collapse during SE, due to its high energy demands and susceptibility to excitotoxicity[63], exhibited preserved mitochondrial gene expression and structural integrity in aKO mice. Similarly, the maintenance of neuronal markers and synaptic proteins in the chronic phase (Day 21) suggests that by limiting the acute inflammatory insult, aKO prevents the maladaptive synapse loss and circuit remodeling that underlie chronic epilepsy and cognitive deficits[64, 65]. The functional preservation of memory and prevention of hyperactive behavior in the chronic phase further validates that the initial suppression of neuroinflammation translates into durable disease modification, supporting the concept that early intervention targeting the neuroinflammatory cascade can prevent epileptogenesis.

## Supporting information

Related to Figure-1

Related to Figure-2

Related to Figure-3

Related to Figure-4

Related to Figure-4

Related to Figure-5

Related to Figure-4

## Summary

Our study redefines the cellular logic of endocannabinoid neuroprotection. We propose that astrocytes serve as the critical reservoir for 2-AG-mediated resilience. By deleting *Mgll* in astrocytes, we effectively uncouple the seizure insult from its devastating inflammatory sequelae. This blockade of glial reactivity preserves the metabolic and structural support systems required for neuronal survival. Consequently, targeting astrocytic endocannabinoid metabolism represents a promising therapeutic strategy to halt the progression of epilepsy and protect against the neuroinflammatory damage of status epilepticus.

## Data availability

Source data are provided with this paper. Additional data supporting the findings of this study are available from the corresponding authors upon request. The single-nucleus RNA sequencing (snRNA-seq) data will be deposited in the NCBI Gene Expression Omnibus for free public access.

## Acknowledgement

This work was supported by startup funds from UT Health San Antonio, Joe R. & Teresa Lozano Long School of Medicine (to C.C.). The authors thank the NIMH Chemical Synthesis and Drug Supply Program for providing JZL184.

## Author contributions

C.Z. and C.C. conceived the project and designed the experiments. C.Z., L.S., M.H., J.L., D.Z., F.G., M.P., J.Z. and C.C. performed the experiments and/or analyzed the data. C.Z. and C.C. wrote the manuscript.

## References

1. Thijs, R.D., et al., Epilepsy in adults. Lancet, 2019. 393(10172): p. 689–701.

2. Vezzani, A., et al., The role of inflammation in epilepsy. Nat Rev Neurol, 2011. 7(1): p. 31–40.

3. Pitkänen, A. and K. Lukasiuk, Mechanisms of epileptogenesis and potential treatment targets. Lancet Neurol, 2011. 10(2): p. 173–86.

4. Zeng, C. and C. Chen, Endocannabinoid signaling in epilepsy. Neurobiol Dis, 2025. 215: p. 107074.

5. Chen, C., Inhibiting degradation of 2-arachidonoylglycerol as a therapeutic strategy for neurodegenerative diseases. Pharmacol Ther, 2023. 244: p. 108394.

6. Castillo, P.E., et al., Endocannabinoid signaling and synaptic function. Neuron, 2012. 76(1): p. 70–81.

7. Sugaya, Y., et al., Crucial Roles of the Endocannabinoid 2-Arachidonoylglycerol in the Suppression of Epileptic Seizures. Cell Rep, 2016. 16(5): p. 1405–1415.

8. Marsicano, G., et al., CB1 cannabinoid receptors and on-demand defense against excitotoxicity. Science, 2003. 302(5642): p. 84–8.

9. Devinsky, O., et al., Cannabidiol in patients with treatment-resistant epilepsy: an open-label interventional trial. Lancet Neurol, 2016. 15(3): p. 270–8.

10. Leuti, A., et al., Bioactive lipids, inflammation and chronic diseases. Adv Drug Deliv Rev, 2020. 159: p. 133–169.

11. Nomura, D.K., et al., Endocannabinoid hydrolysis generates brain prostaglandins that promote neuroinflammation. Science (New York, N.Y.). 334(6057): p. 809–13.

12. Lu, Y., et al., Endocannabinoid 2-arachidonylglycerol protects primary cultured neurons against LPS-induced impairments in rat caudate nucleus. J Mol Neurosci, 2014. 54(1): p. 49–58.

13. von Rüden, E.L., et al., Inhibition of monoacylglycerol lipase mediates a cannabinoid 1-receptor dependent delay of kindling progression in mice. Neurobiol Dis, 2015. 77: p. 238–45.

14. Zareie, P., et al., Anticonvulsive effects of endocannabinoids; an investigation to determine the role of regulatory components of endocannabinoid metabolism in the Pentylenetetrazol induced tonic-clonic seizures. Metab Brain Dis, 2018. 33(3): p. 939–948.

15. Terrone, G., et al., Inhibition of monoacylglycerol lipase terminates diazepam-resistant status epilepticus in mice and its effects are potentiated by a ketogenic diet. Epilepsia, 2018. 59(1): p. 79–91.

16. Hu, M., et al., Enhancing endocannabinoid signalling in astrocytes promotes recovery from traumatic brain injury. Brain, 2022. 145(1): p. 179–193.

17. Chen, R., et al., Monoacylglycerol lipase is a therapeutic target for Alzheimer’s disease. Cell Rep, 2012. 2(5): p. 1329–39.

18. Du, H., et al., Inhibition of COX-2 expression by endocannabinoid 2-arachidonoylglycerol is mediated via PPAR-γ. Br J Pharmacol, 2011. 163(7): p. 1533–49.

19. Zhang, J., et al., Synaptic and cognitive improvements by inhibition of 2-AG metabolism are through upregulation of microRNA-188-3p in a mouse model of Alzheimer’s disease. J Neurosci, 2014. 34(45): p. 14919–33.

20. Song, Y., et al., A novel mechanism of synaptic and cognitive impairments mediated via microRNA-30b in Alzheimer’s disease. EBioMedicine, 2019. 39: p. 409–421.

21. Gao, F., et al., TDP-43 drives synaptic and cognitive deterioration following traumatic brain injury. Acta Neuropathol, 2022. 144(2): p. 187–210.

22. Zhang, J., et al., A Combination of Low-Dose Δ(9)-THC and Celecoxib as a Therapeutic Strategy for Alzheimer’s Disease. Aging Dis, 2025.

23. Zhang, J., et al., Inhibition of monoacylglycerol lipase prevents chronic traumatic encephalopathy-like neuropathology in a mouse model of repetitive mild closed head injury. J Cereb Blood Flow Metab, 2015. 35(3): p. 443–53.

24. Sang, N., et al., Postsynaptically synthesized prostaglandin E2 (PGE2) modulates hippocampal synaptic transmission via a presynaptic PGE2 EP2 receptor. J Neurosci, 2005. 25(43): p. 9858–70.

25. Zhang, J. and C. Chen, Endocannabinoid 2-arachidonoylglycerol protects neurons by limiting COX-2 elevation. J Biol Chem, 2008. 283(33): p. 22601–11.

26. Zhang, J., et al., A Combination of Low-Dose Δ ^9^-THC and Celecoxib as a Therapeutic Strategy for Alzheimer’s Disease. Aging and disease, 2025: p. 0-.

27. Sun, L., et al., Selective Inactivation of Astrocytic Monoacylglycerol Lipase for Alzheimer’s Disease Therapy. bioRxiv, 2025.

28. Zhu, D., et al., Overabundant endocannabinoids in neurons are detrimental to cognitive function. bioRxiv, 2024.

29. Germain, P.L., et al., Doublet identification in single-cell sequencing data using scDblFinder. F1000Res, 2021. 10: p. 979.

30. Zhu, D., et al., Inhibition of 2-arachidonoylglycerol degradation enhances glial immunity by single-cell transcriptomic analysis. J Neuroinflammation, 2023. 20(1): p. 17.

31. Jin, S., M.V. Plikus, and Q. Nie, CellChat for systematic analysis of cell–cell communication from single-cell transcriptomics. Nature Protocols, 2025. 20(1): p. 180–219.

32. Alvarez, M.J., et al., Functional characterization of somatic mutations in cancer using network-based inference of protein activity. Nature Genetics, 2016. 48(8): p. 838–847.

33. Türei, D., et al., Integrated intra- and intercellular signaling knowledge for multicellular omics analysis. Molecular Systems Biology, 2021. 17(3): p. MSB20209923.

34. Müller-Dott, S., et al., Expanding the coverage of regulons from high-confidence prior knowledge for accurate estimation of transcription factor activities. Nucleic Acids Research, 2023. 51(20): p. 10934–10949.

35. Mardones, M.D. and K. Gupta, Transcriptome Profiling of the Hippocampal Seizure Network Implicates a Role for Wnt Signaling during Epileptogenesis in a Mouse Model of Temporal Lobe Epilepsy. Int J Mol Sci, 2022. 23(19).

36. Wallace, M.J., et al., The endogenous cannabinoid system regulates seizure frequency and duration in a model of temporal lobe epilepsy. J Pharmacol Exp Ther, 2003. 307(1): p. 129–37.

37. Fezza, F., et al., Distinct modulation of the endocannabinoid system upon kainic acid-induced in vivo seizures and in vitro epileptiform bursting. Mol Cell Neurosci, 2014. 62: p. 1–9.

38. Chen, C., Homeostatic regulation of brain functions by endocannabinoid signaling. Neural Regen Res, 2015. 10(5): p. 691–2.

39. Chali, F., et al., Lipid markers and related transcripts during excitotoxic neurodegeneration in kainate-treated mice. Eur J Neurosci, 2019. 50(1): p. 1759–1778.

40. Du, H., et al., Inhibition of COX-2 expression by endocannabinoid 2-arachidonoylglycerol is mediated via PPAR-gamma. Br J Pharmacol, 2011. 163(7): p. 1533–49.

41. Kow, R.L., et al., Modulation of pilocarpine-induced seizures by cannabinoid receptor 1. PLoS One, 2014. 9(4): p. e95922.

42. Noriega-Prieto, J.A., et al., Distinct endocannabinoids specifically signal to astrocytes or neurons in the adult mouse hippocampus. Nature Neuroscience, 2025.

43. Viader, A., et al., Metabolic Interplay between Astrocytes and Neurons Regulates Endocannabinoid Action. Cell Rep, 2015. 12(5): p. 798–808.

44. Guggenhuber, S., et al., Impaired 2-AG Signaling in Hippocampal Glutamatergic Neurons: Aggravation of Anxiety-Like Behavior and Unaltered Seizure Susceptibility. Int J Neuropsychopharmacol, 2015. 19(2).

45. Terrone, G., A. Salamone, and A. Vezzani, Inflammation and Epilepsy: Preclinical Findings and Potential Clinical Translation. Curr Pharm Des, 2017. 23(37): p. 5569–5576.

46. Sano, F., et al., Reactive astrocyte-driven epileptogenesis is induced by microglia initially activated following status epilepticus. JCI Insight, 2021. 6(9).

47. Kong, S., et al., Cell-specific NFIA upregulation promotes epileptogenesis by TRPV4-mediated astrocyte reactivity. J Neuroinflammation, 2023. 20(1): p. 247.

48. Vezzani, A., et al., Astrocytes in the initiation and progression of epilepsy. Nat Rev Neurol, 2022. 18(12): p. 707–722.

49. Covelo, A., et al., CB1R-dependent regulation of astrocyte physiology and astrocyte-neuron interactions. Neuropharmacology, 2021. 195: p. 108678.

50. Beggiato, S., et al., Astrocytic palmitoylethanolamide pre-exposure exerts neuroprotective effects in astrocyte-neuron co-cultures from a triple transgenic mouse model of Alzheimer’s disease. Life Sci, 2020. 257: p. 118037.

51. Microglia eliminate inhibitory synapses and drive neuronal hyperexcitability in epilepsy. Nature Neuroscience, 2025. 28(7): p. 1370–1371.

52. Bolaños, J.P., Bioenergetics and redox adaptations of astrocytes to neuronal activity. J Neurochem, 2016. 139 Suppl 2(Suppl Suppl 2): p. 115–125.

53. Jiwaji, Z., et al., Reactive astrocytes acquire neuroprotective as well as deleterious signatures in response to Tau and Aß pathology. Nature Communications, 2022. 13(1): p. 135.

54. Jimenez-Blasco, D., et al., Glucose metabolism links astroglial mitochondria to cannabinoid effects. Nature, 2020. 583(7817): p. 603–608.

55. Mallach, A., et al., Microglia-astrocyte crosstalk in the amyloid plaque niche of an Alzheimer’s disease mouse model, as revealed by spatial transcriptomics. Cell Reports, 2024. 43(6): p. 114216.

56. Lee, H.-G., et al., Neuroinflammation: An astrocyte perspective. Science Translational Medicine. 15(721): p. eadi7828.

57. Wilson, R.I., G. Kunos, and R.A. Nicoll, Presynaptic Specificity of Endocannabinoid Signaling in the Hippocampus. Neuron, 2001. 31(3): p. 453–462.

58. Eldeeb, K., S. Leone-Kabler, and A.C. Howlett, CB1 cannabinoid receptor-mediated increases in cyclic AMP accumulation are correlated with reduced Gi/o function. J Basic Clin Physiol Pharmacol, 2016. 27(3): p. 311–22.

59. Lehrke, M. and M.A. Lazar, The many faces of PPARgamma. Cell, 2005. 123(6): p. 993–9.

60. Grabner, G.F., et al., Deletion of Monoglyceride Lipase in Astrocytes Attenuates Lipopolysaccharide-induced Neuroinflammation. J Biol Chem, 2016. 291(2): p. 913–23.

61. Rockwell, C.E., et al., Interleukin-2 suppression by 2-arachidonyl glycerol is mediated through peroxisome proliferator-activated receptor gamma independently of cannabinoid receptors 1 and 2. Mol Pharmacol, 2006. 70(1): p. 101–11.

62. Senn, L., et al., Is the peroxisome proliferator-activated receptor gamma a putative target for epilepsy treatment? Current evidence and future perspectives. Pharmacology & Therapeutics, 2023. 241: p. 108316.

63. Bartsch, T., et al., Selective neuronal vulnerability of human hippocampal CA1 neurons: lesion evolution, temporal course, and pattern of hippocampal damage in diffusion-weighted MR imaging. J Cereb Blood Flow Metab, 2015. 35(11): p. 1836–45.

64. Wang, Q., et al., The Progress of Cognitive Dysfunction Impairment Caused by Temporal Lobe Epilepsy. J Mol Neurosci, 2025. 75(3): p. 81.

65. Xiao, L., et al., Association of synaptic density and cognitive performance in temporal lobe epilepsy: Humans and animals PET imaging study with [18F]SynVesT-1. Psychiatry and Clinical Neurosciences, 2024. 78(8): p. 456–467.

