## Supplementary material for "Inhibition of Endocannabinoid Degradation in Astrocytes Reprograms Glial Reactivity and Prevents Seizure Sequelae": Related to Figure-1

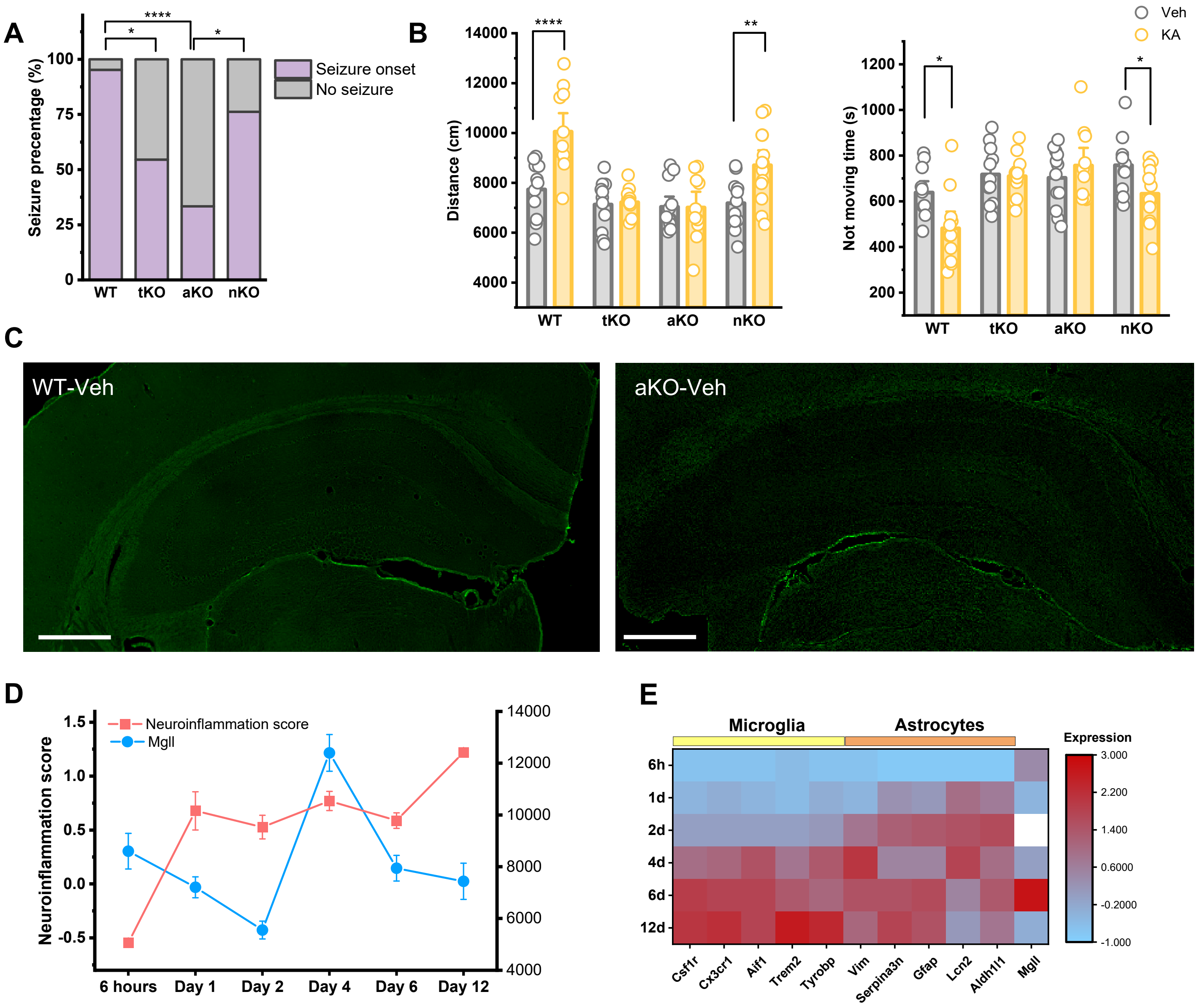


**Figure S1. Astrocyte-specific Mgll deletion attenuates kainic acid (KA)-induced seizure susceptibility and hippocampal neuroinflammation** (A) Quantification of the percentage of mice developing seizures in WT, TKO, AKO, and NKO mice in one hour (WT: 20 mice, TKO: 22 mice, AKO: 25 mice, NKO: 21 mice). Statistical significance was determined by a Chi-Squared Test with Bonferroni multiple test, *$p < 0.05$,****$p < 0.001$. (B) Quantification of hyperactive and anxiety-like behaviors in the Open Field test, showing total distance traveled and normalized freezing time of both the baseline and post-seizure injury. Data are means ± SEM. *$p < 0.05$,**$p < 0.01$, ***$p < 0.005$, ****$p < 0.001$ (n=11-13, t-test in each genotype). (C) Representative immunofluorescence images of Fluoro-Jade C (FJC) stainingin the hippocampus of WT and AKO mice following vehicle treatment. Scale bars: 500 µm. (D) Temporal profile of the Neuroinflammation score (red) and Mgll expression (blue) over 14 days in CA1. (E) Heatmap illustrating the temporal mRNA expression profiles of *Mgll*, astrocytic and microglial makrers in the hippocampus of mice treated with saline or KA at 6 hours, days 1, 2,4, 6, and 12 post-injection. The color scale indicates relative expression levels **.**
