## Supplementary material for "Inhibition of Endocannabinoid Degradation in Astrocytes Reprograms Glial Reactivity and Prevents Seizure Sequelae": Related to Figure-2

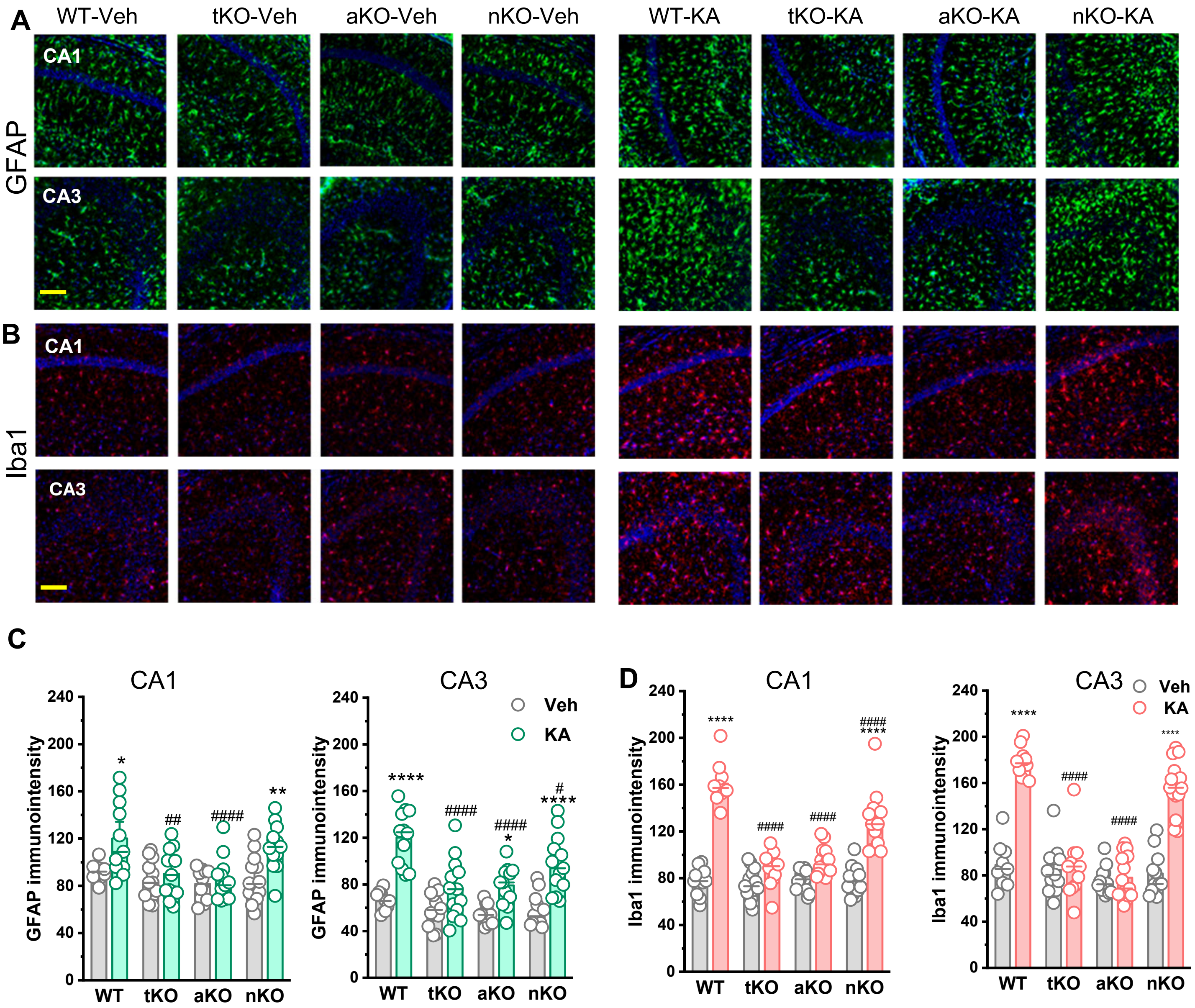


**Figure S2. Astrocyte-specific Mgll deletion attenuates kainic acid (KA)-induced gliosis.** (A-B) Representative immunofluorescence images of GFAP (F, green) and Iba1 (G, red) staining in the CA1, and CA3 regions of the hippocampus across different genotypes and treatments. Nuclei are stained with DAPI (blue). Scale bar: 100 µm. (C–D) Quantification of GFAP (C) and Iba1 (D) immunofluorescence intensity in the CA1 and CA3 regions (n=4-5 mice). Data represent individual slice values (2–3 slices per mouse) from n = 4 mice per group. Data are means ± SEM. *p < 0.05,**p < 0.01, ****p < 0.001, compared with Veh in each genotype, #p < 0.01, ##p < 0.01, ####p < 0.001 compared with WT-KA (ANOVA with Turkey test).
