## Supplementary material for "Inhibition of Endocannabinoid Degradation in Astrocytes Reprograms Glial Reactivity and Prevents Seizure Sequelae": Related to Figure-3

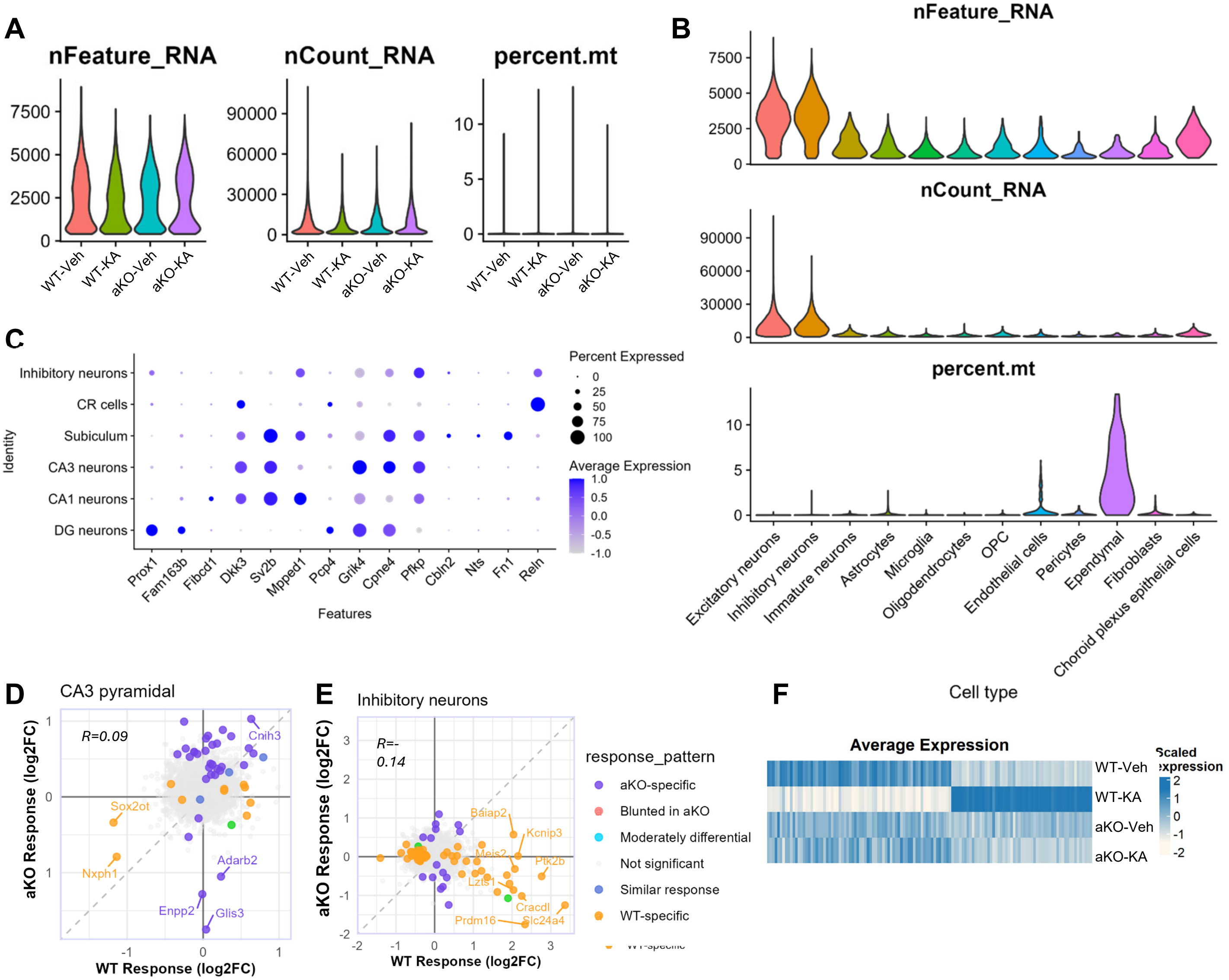


**Figure S3. Single-nucleus RNA sequencing reveals distinct cell-type-specific transcriptional responses to KA in AKO mice**. (A) Quality control (QC) metrics by experimental group. Violin plots showing the distribution of the number of unique genes (nFeature_RNA), total RNA molecules (nCount_RNA), and the percentage of mitochondrial reads (percent.mt) for the four groups: WT-Veh, WT-KA, aKO-Veh, aKO-KA. The distributions remain consistent across groups, indicating no significant technical bias introduced by the treatment or genotype. (B) QC metrics by annotated cell type. Violin plots of nFeature_RNA, nCount_RNA, and percent.mt categorized by major hippocampal cell populations. Note the higher mitochondrial content in ependymal cells and higher feature counts in neuronal populations, consistent with typical single-nucleus RNA-seq profiles.(C) Dot plot of canonical neuronal cell-type markers.(D-E) Scatter plots comparing the transcriptional response to KA (log2 Fold Change) between WT (x-axis) and aKO (y-axis) mice in CA3 pyramidal cells (D) and inhibitory neurons. (F) Heatmaps showing the relative expression of differentially expressed genes (DEGs) across experimental groups in Inhibitory granular cells.
