## Supplementary material for "Inhibition of Endocannabinoid Degradation in Astrocytes Reprograms Glial Reactivity and Prevents Seizure Sequelae": Related to Figure-4

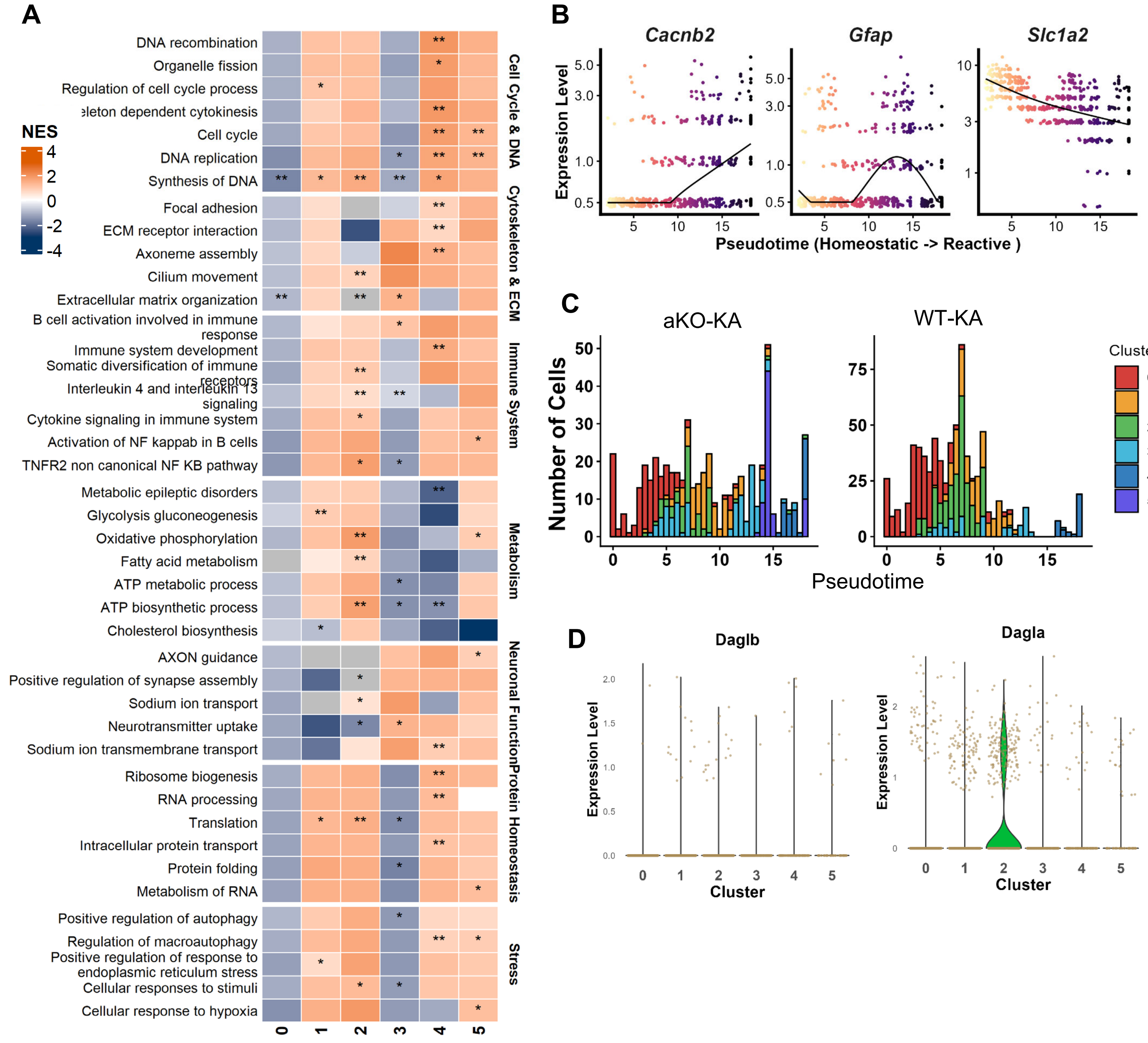


**Figure S4. Mgll deletion alters astrocyte functional states.**(A) Heatmap displaying the Normalized Enrichment Score (NES) for various biological processes across clusters 0–5 via Gene Set Enrichment Analysis (GSEA). Stars indicate statistical significance (* $p < 0.05$, ** $p < 0.01$). (B) Expression levels of *Cacnb2*, *Gfap*, and *Slc1a2* plotted against a pseudotime trajectory representing the transition from homeostatic to reactive astrocyte states.(C) Histograms showing the number of cells across pseudotime for aKO-KA and WT-KA groups, colored by cluster identity. The aKO-KA group displays a higher density of cells in cluster 2 compared to the WT-KA group. (D) Violin plots showing the expression levels of endocannabinoid synthesizing enzymes *Daglb* and *Dagla* across clusters 0–5. *Dagla* shows enriched expression specifically in Cluster 2.
