## Supplementary material for "Inhibition of Endocannabinoid Degradation in Astrocytes Reprograms Glial Reactivity and Prevents Seizure Sequelae": Related to Figure-4

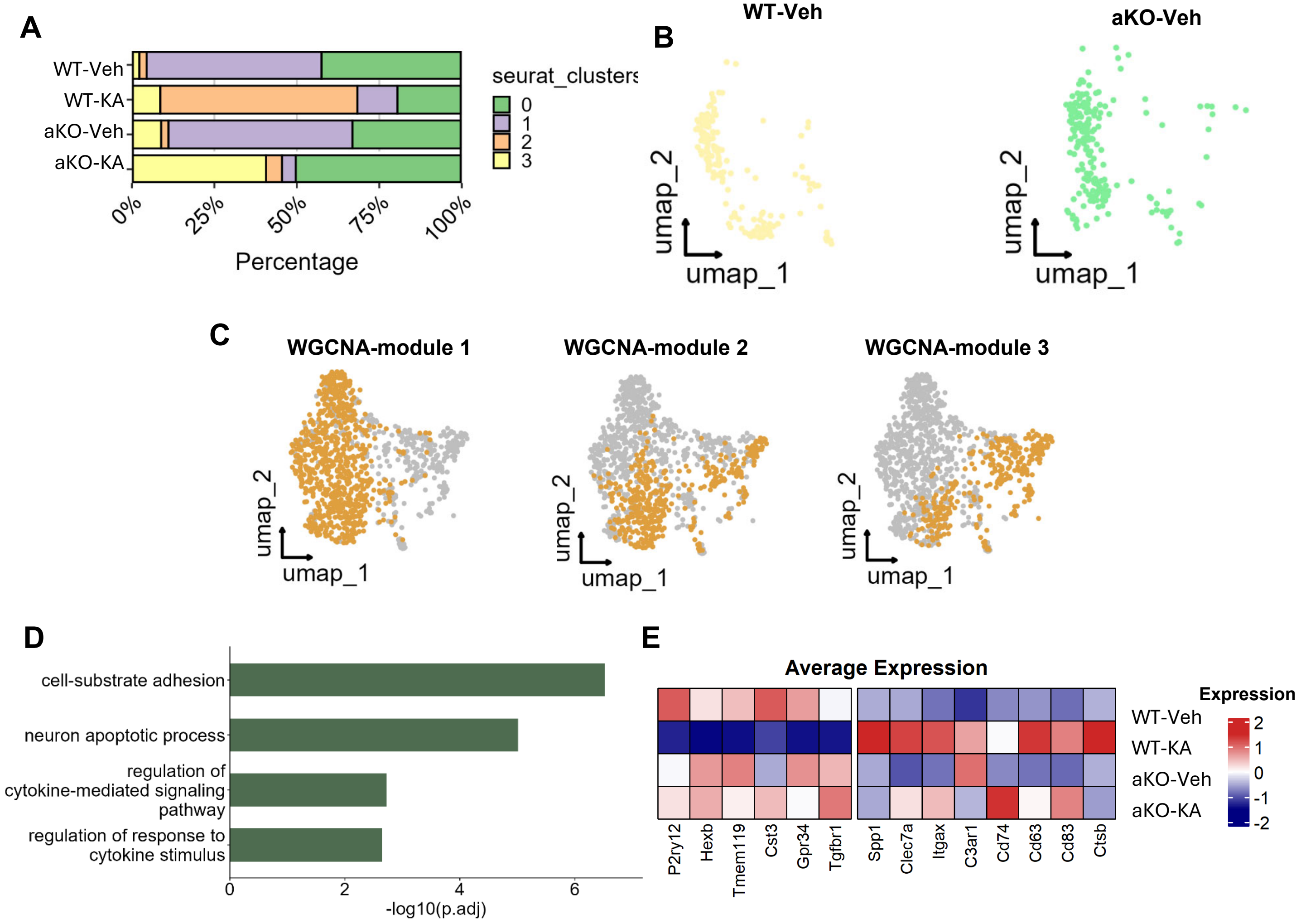


**Figure S5. Mgll deletion suppresses disease-associated microglia (DAM) activation**. (A) Bar plot showing the percentage of cells from each cluster (0–5) contributing to the four experimental groups. (B) Dimensional reduction plots showing the distribution of astrocytes in wt_veh and ako_veh groups. (C) UMAP plots highlighting the expression density of three specific gene modules (M1, M2, and M3) from WGCNA across the astrocyte population. (D) Bar chart showing significant GO terms for hub genes of WGCNA M3 module. (E) Heatmap showing scaled average expression of homeostatic markers (e.g., P2ry12, Tmem119) and reactive/phagocytic markers (e.g., Spp1, Cd74) across groups.
