## Supplementary material for "Inhibition of Endocannabinoid Degradation in Astrocytes Reprograms Glial Reactivity and Prevents Seizure Sequelae": Related to Figure-5

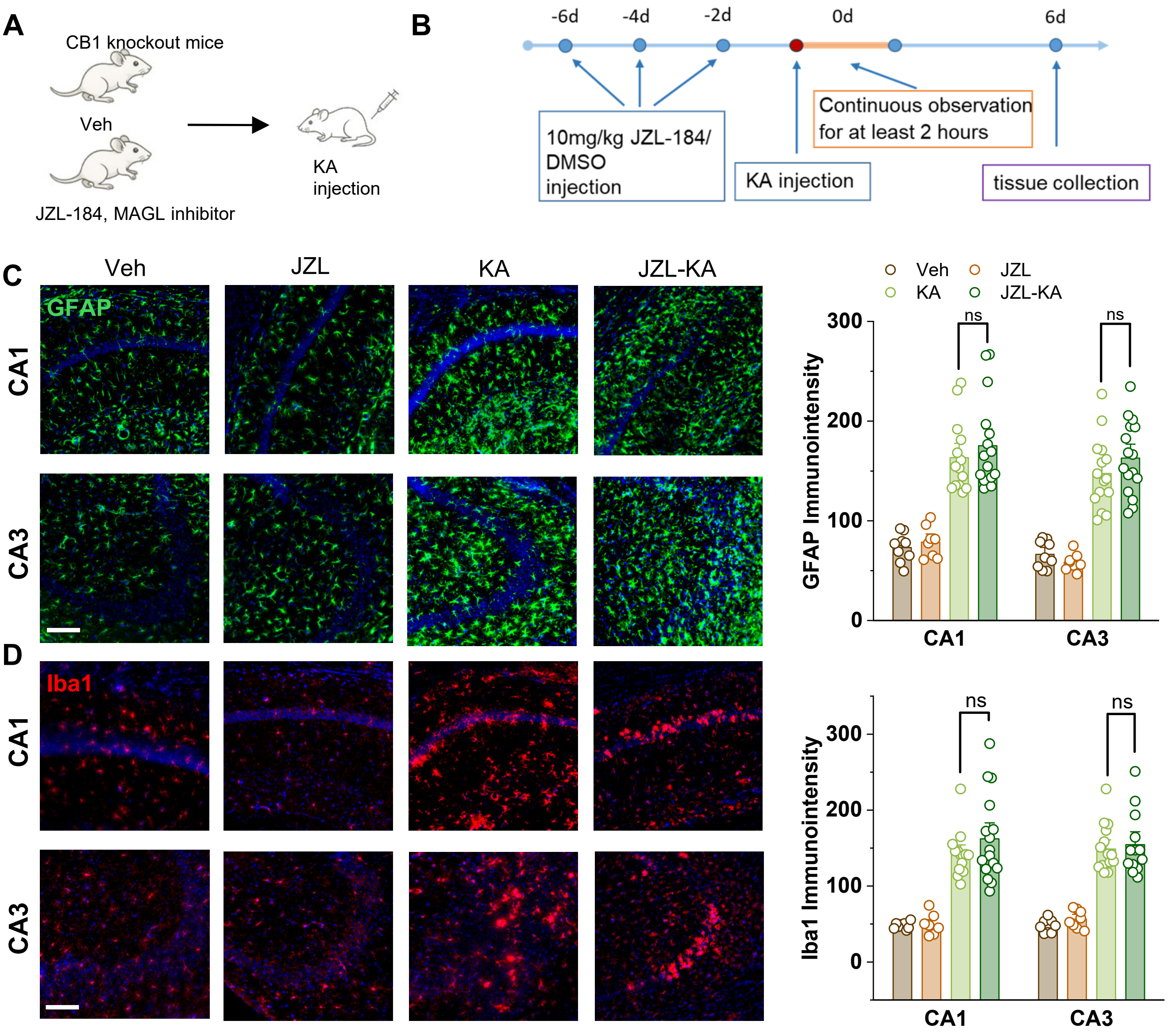


**Figure S6. The neuroprotective effect of MAGL inhibition is dependent on CB1 receptor signaling and mediated by PPAR-γ activation.** (A-B) Schematic of the experimental design (A) and timeline (B) for JZL-184 or DMSO (vehicle) administration followed by KA-induced epilepsy in Cnr1$^{-/-}$ mice. (C) Representative images of GFAP (green) staining in the CA1 and CA3 hippocampal regions with quantification. Scale bar: 100 μm. Data are means ± SEM. Data represent individual slice values (3–4 slices per mouse) from 4 mice in KA groups and 3 mice in Veh groups . ns $p > 0.05$ (ANOVA with Turkey test). (D) Representative images of Iba1 (red) staining in the CA1 and CA3 hippocampal regions with quantification. Scale bar: 100 μm. Data are means ± SEM. Data represent individual slice values (3–4 slices per mouse) from 4 mice in KA groups and 3 mice in Veh groups . ns $p > 0.05$ (ANOVA with Turkey test).
